# Neural population dynamics of direct electrical stimulation of neocortex

**DOI:** 10.1101/2025.10.28.685195

**Authors:** Jordan L. Hickman, Grant Hughes, Eashan Sahai, Moriah Miles, Daniel J. Denman

## Abstract

Intracranial electrical stimulation is foundational in neuroscience and clinical neuromodulation, yet how applied electrical fields engage neural tissue and influences perception remains unclear. Using three orthogonal Neuropixels probes in mouse visual areas, we recorded extracellular voltages and spikes with sub-millisecond resolution in three dimensions around a stimulation source. The evoked potential extended asymmetrically and grew sub-linearly with amplitude, influenced by anatomical heterogeneity. Increasing amplitude increased the density, but not spatial extent, of directly responsive neurons, but fewer than 5% of neurons within the evoked potential were activated. Fast-spiking interneurons were recruited nearer the source, whereas pyramidal neurons showed anisotropic activation. Direct responses were polarity- and amplitude-selective. Sparse direct spiking drove large, asymmetric cortical responses and modulated >50% of neurons at higher amplitudes. Perceptually, mice weakly detected single pulses, and performance did not improve with amplitude. Thus, anatomy and spatiotemporal patterning—not spike counts—govern physiological and perceptual impact, informing biomimetic, multi-contact prostheses.

## INTRODUCTION

Intracranial electrical stimulation is a powerful tool for modulating neural activity, with widespread applications in basic neuroscience to assess neural function and behavior [1–3] and clinical medicine. Clinically, intracranial electrical stimulation is used to treat movement disorders including Parkinson’s [4–6], essential tremor [7], and dystonia [8]; disorders of mental health including depression [9], obsessive-compulsive disorder [10], and schizophrenia [11]; and epilepsy [12]. Beyond these indications for therapeutic neuromodulation, there is also growing interest in using electrical stimulation in cortical prostheses, devices that aim to restore lost perception by encoding information through stimulating the brain [13–22].

Electrical stimulation has long been used as a tool to interrogate and restore sensory function. Early work demonstrated that direct electrical stimulation of the human visual cortex can evoke rudimentary percepts termed “phosphenes” [23], laying the groundwork for further exploration of cortical visual prostheses [24–28]. Similarly, electrical stimulation in the somatosensory cortex elicits localized tactile percepts [15], suggesting feasibility for somatosensory neuroprosthetics and for providing sensory feedback to improve motor BCIs [14]. However, a major limitation of sensory cortical prosthetics to date is the inability to generate percepts that closely resemble natural sensory experiences [20, 29]. Increasing the number of stimulating electrodes has expanded coverage of retinotopic and somatotopic space, enabling perception of a greater number of stimuli, and facilitated simultaneous stimulation at multiple sites to create additive percepts, yet the quality of percepts remains limited [18, 20, 29, 30].

Generally, approaches to enhancing the quality of electrically-evoked percepts seek to leverage spatial and temporal interactions of multiple electrical stimuli. For example, exploiting putative spatial specificity to isolate laminar structure, and stimulating from multiple sites within functional column, can improve detection thresholds in rodents [31, 32]. Others focus on refining stimulation paradigms to temporally mimic sensory stimuli: dynamic current steering in the primary visual cortex of sighted and blind humans can evoke percepts of letter forms, whereas static stimulation is unable to [33]. In the somatosensory cortex, spatially and temporally structured pulse trains that mimic structured spike trains derived from natural touch have been shown to evoke percepts that more closely resemble real tactile experiences [34, 35]. While these stimulation approaches can improve the perceptual quality and the number of discriminable features, they remain far from recreating naturalistic stimuli. A major limitation in the design of these spatiotemporal stimulation paradigms is the incomplete empirical understanding of the direct impact of electrical stimulation on intact neural tissue.

Direct measurements of the evoked fields and neural response to electrical stimulation have been technically challenging due to electrical artifacts in electrophysiological recordings, spatial limits of electrophysiology techniques, temporal imprecision in imaging approaches, and ambiguity in the transfer function relating voltage or calcium changes to the recorded fluorescence. Still, studies have long sought to characterize the neural response to electrical stimulation [36]. Classic work by Stoney et al. in cat motor cortex demonstrated that electrical stimulation excites neurons within a roughly spherical volume that expands with increasing current intensity by using pyramidal tract activation as a proxy for the affected volume [37]. This led to the hypothesis that higher currents recruit neurons at increasing distances. Four decades later, Histed et al. revisited this with two-photon calcium imaging and arrived at a different picture. They found that even low-current electrical stimulation (5-10 μA) sparsely activates neurons distributed hundreds of microns to millimeters away from the electrode. These activated cells were scattered in space, and their pattern changed with small (tens of μm) electrode movements, suggesting that the volume of affected tissue is small and stimulation primarily triggers local axonal and dendritic processes rather than cell bodies [38]. These studies, using distinct approaches, framed two contrasting views: one of an expanding volume of uniform tissue activation with higher currents [37] and another of a fixed local stimulation zone yielding widespread but sparse neuronal activation patterns[38]. Computational modeling studies have attempted to reconcile these differences, arguing the spatial distribution of activated somata remains relatively constant, with higher currents mainly increasing the density of activated neurons with that area [39]; this hypothesis remains to be empirically verified.

Here, we aim to measure the spatial and temporal extents of the effects electrical stimulation on underlying neural tissue – the induced voltage changes and the resulting responses of populations of single neurons. To do so, we use multiple spatially registered high-density Neuropixels electrophysiological devices to capture electrically-evoked activity at fine spatial (10 µm) and temporal (sub-millisecond) resolution, varying intracranial stimulation amplitude and polarity of single pulses. With these data, we quantify the spatial properties of stimulation-evoked neural activity, how single neurons directly respond to stimulation, and how single-neuron direct responses propagate into circuit-wide activity. These measures allow us to validate models of brain conductivity and electrical stimulation specificity for all uses of intracranial electrical stimulation that seek to modulate neural dynamics, including both deep brain stimulation and cortical prostheses. Finally, we evaluate the relationship between induced neural dynamics and perception using matched electrical stimulation in a visual detection test paradigm.

## RESULTS

### Orthogonal multi-Neuropixel recordings for spatial measurement of electrical stimulation in awake mice

To measure the effects of electrical stimulation in neocortex, we simultaneously inserted three Neuropixels 1.0 [40], orthogonally to each other, to measure neural activity in three dimensions around a stimulation source (a 16-channel stimulating electrode, see Methods) within the primary visual cortex in awake, head-fixed mice (Figure 1a). This configuration allowed continuous sampling within and across multiple brain regions, including the primary visual cortex (V1), higher-order visual areas such as anteromedial (AM), lateral (LM), posterolateral (PL), as well as sub-neocortical regions, across a white matter tract, including hippocampus (CA1) and subiculum (Figure 1c). The corpus callosum – a dense sheet of white matter – separates cortical and subcortical structures and provides a distinct anatomical reference point to quantify the influence of anatomical heterogeneity on the spatial spread of electrical stimulation. We targeted stimulation to layer 5 of V1 due to its role in cortical output pathways, dense activity, and anatomical proximity to the white matter boundary (Figure 1i). Each recording Neuropixels probe and the stimulating electrode array were registered in mouse Common Coordinate Framework (CCF) brain coordinate space [41] to measure anatomical and distance effects of stimulation responses and allow comparisons across recordings. To align each probe and the stimulating electrode accurately, we combined an *a priori* stereotactic plan based on fixed probe geometry, where each probe was positioned at consistent angles and relative distances, with post-hoc verification using histology and electrophysiological landmarks (Figures 1a–1e, Methods). This alignment assigns each Neuropixels channel and stimulating electrode contact a CCF coordinate; these coordinates are mapped to anatomical brain regions and also allow 3-dimensional spatial measurements across recordings (Figures 1f–1g).

**Figure 1.**
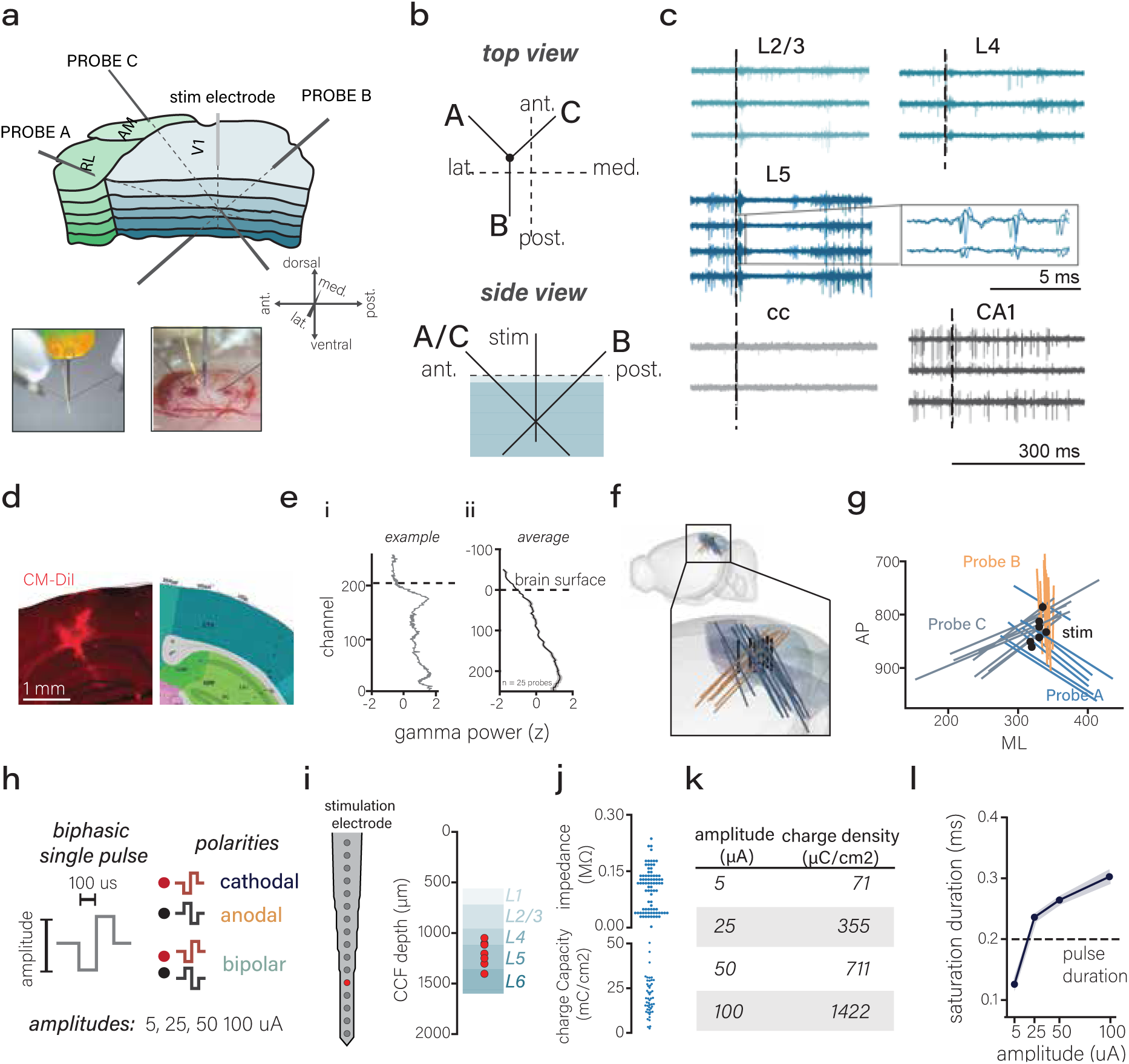
Orthogonal multi-neuropixel recordings for spatial measurement of electrical stimulation in awake mice. **(a)** Schematic of orthogonal probe geometry in brain space. Bottom-left inset: photo of three Neuropixels probes and one stimulating electrode arranged at full-insertion depth. Bottom-right inset: photo of probe geometry during insertion through a small craniotomy. **(b)** Top-down (upper panel) and lateral (lower panel) schematic views of orthogonal probe geometry. **(c)** Example neural response to 50 µA bipolar electrical stimulation grouped by brain region for 300 ms post-stimulation. Inset: 10 ms window highlighting stimulation-evoked spiking activity in Layer 5 neurons. **(d)** Histological alignment of probe positions using fluorescent dye labeling (CM-DiI). **(e)** Electrophysiological alignment example (i) and average (ii), using LFP gamma-band power to identify cortical surface and probe depth (mean and SEM error, N = 9 recordings, n = 25 probes). **(f)** Three-dimensional rendering of Neuropixels probes (n = 25 probes from N = 9 recordings) registered into Allen Common Coordinate Framework (CCF) brain space. **(g)** Two-dimensional (anterior-posterior vs. medial-lateral) representation of probe geometry for all recordings (n = 25 probes, N = 9 recordings) in CCF brain space. **(h)** Single-pulse electrical stimulation parameters illustrating variations in amplitude and polarities used in experiments. **(i)** Schematic of the 16-channel stimulating electrode (left panel), with summary of active contact depth positions across experiments (right panel, N = 9 recordings). **(j)** Average contact impedance (top) and charge capacity (bottom) for stimulation contacts. **(k)** Table converting electrical stimulation amplitudes (µA) to corresponding charge densities (µC/cm²). **(l)** Duration of recording amplifier duration by amplitude (mean and SEM error bars, One-way ANOVA, F = 105.6, p < 0.001).

During each recording session, we systematically varied electrical stimulation parameters of single constant-current electrical pulses, focusing on current amplitude and polarity (Figure 1h, Methods). We used a linear 16-channel stimulating electrode (Neuronexus, A1x16-5mm-50-703) with iridium oxide site activation to decrease contact impedance and increase charge delivery for more efficacious stimulation delivery. The average contact impedance was 0.10*±*0.06 MΩ (Figure 1j, top), and the average contact charge capacity was 25.90 *±*9.69 mC/cm² (Figure 1j, bottom). We describe the charge magnitude in amperes (µA) but include a conversion to charge density (Figure 1k), allowing findings to be better extrapolated to other electrode configurations. Simultaneous electrophysiology and electrical stimulation – especially with recording electrodes in close proximity to the stimulation source – can result in saturating electrical artifacts that obscure neural data. In our recordings, we were able to exploit Neuropixels 1.0 band-specific amplifier gain and the rapid settling of Neuropixels amplifiers to observe the neural response almost immediately following stimulation. On average, following a 0.2 ms total duration electrical pulse, saturation persists for approximately 0.14 ± 0.00 ms at 5 µA, 0.24 ± 0.01 ms at 25 µA, 0.26 ± 0.01 ms at 50 µA, and 0.30 ± 0.01 ms at 100 µA (Figure 1l, One-way ANOVA, F = 105.62, p < 0.001). In rare cases, saturation at channels extremely close to the stimulating electrode extends up until 1 ms for cathodal-leading monopolar and cathodal-leading bipolar (Figure S1a, i-iii). Overall we were able to observe neural activation across polarities and amplitudes (Figure S1b) despite the increases in saturation with amplitude (2-way ANOVA, F = 236.04, p < 0.001) and polarity alone did not significantly change saturation duration (F = 2.48, p = 0.09).

### Evoked potential spatial extent increases sub-linearly with amplitude

We first sought to characterize the spatial spread of the electric field generated by a single electrode or a pair of adjacent electrodes during electrical stimulation. In an idealized scenario of a homogeneous, isotropic medium, the electric field from a point current source can be approximated by Gauss’s law:

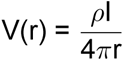

where *ρ* is the tissue resistivity (Ω *·* m), I is the injected current (in amperes), and r is the radial distance from the source (in meters) [42]. Applications of Gauss’s law to model of brain stimulation assume uniform brain conductivity, neglecting potential anatomical and conductive variations such as differences in axonal myelination, dendritic branching, and somatic size. Consequently, these models predict isotropic spatial boundaries for evoked potentials that increase linearly with current amplitude [39, 42, 43]. Local anatomical structures and heterogeneous tissue conductivity could significantly influence both the magnitude and spatial profile of electrically evoked potentials (Figure 2d), but direct measurements of this have been elusive. We first employed the idealized point-source model with a uniform cerebral cortex resistivity of 5.8 Ω *·* m [44, 45] to calculate expected voltage distributions and spatial extents at experimental currents (5, 25, 50, 100 µA) (Figures 2a–2c, Methods). These theoretical predictions provided a null model for symmetry and extent by amplitude to compare to *in vivo* observations

**Figure 2.**
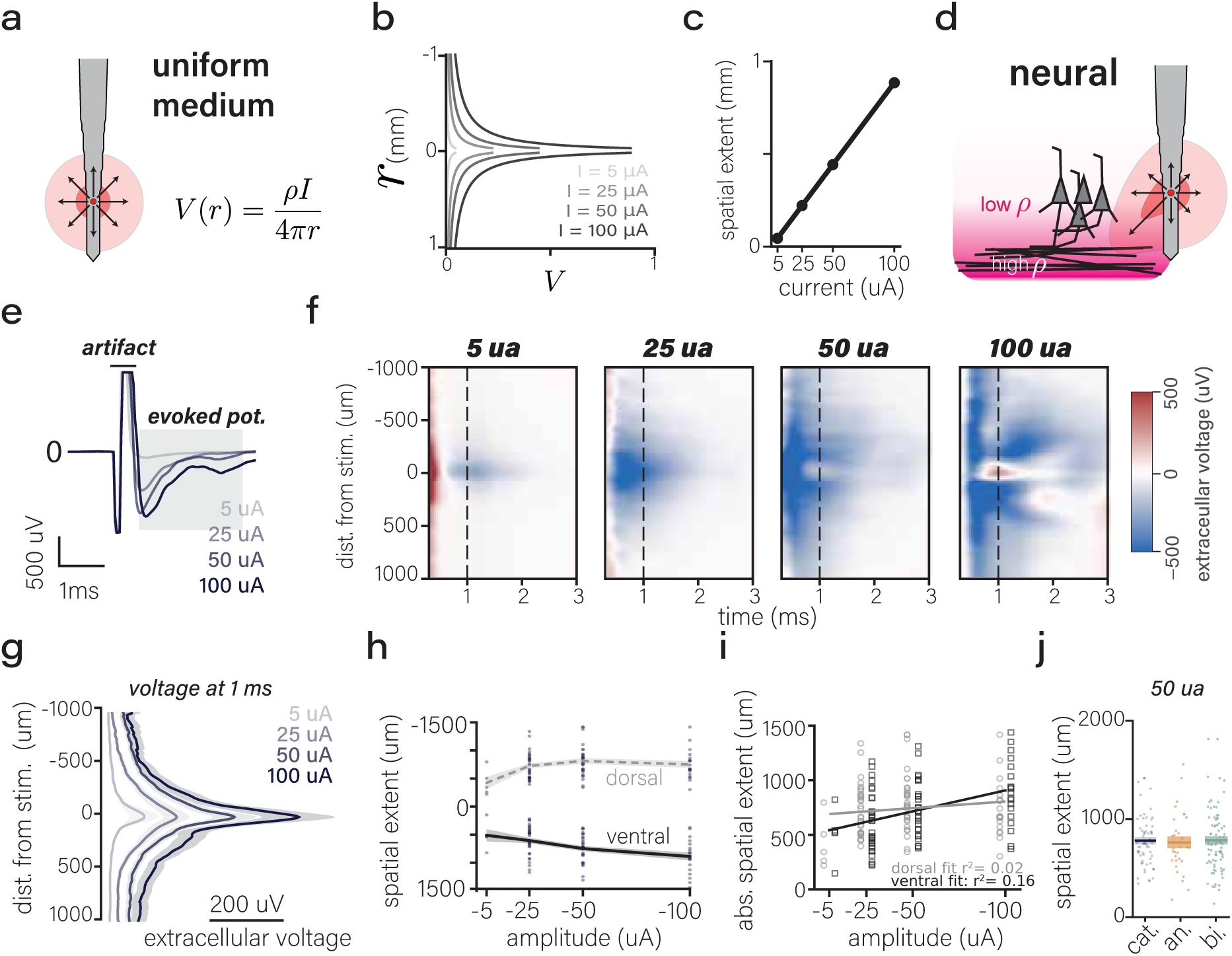
Evoked potential spatial extent increases sub-linearly with amplitude. **(a)** Schematic representation of electrical stimulation as a point source in a homogeneous resistivity environment according to Gauss’ equation (inset) where rho is cortex resistivity (5.56 Ohm*M), r is the radius from the stimulation source, V is the electric potential (volts), and I is the current (amperes). **(b)** Theoretical electric potential predicted by the point-source model at increasing radii (excluding the first 50 µm around stimulation site) for stimulation amplitudes of 5, 25, 50, and 100 µA. **(c)** The radial spatial extent where voltage returns to zero as a function of current amplitude. **(d)** Schematic illustrating inhomogeneity of conductivities of white matter and gray matter. **(e)** Mean extracellular potential from a representative single Neuropixel channel in the AP band demonstrating both artifact and subsequent evoked potential to 5, 25, 50, and 100 µA cathodal single pulses (N = 9 recordings, n = 25 probes, mean and SEM error clouds). **(f)** Heatmaps showing mean extracellular potentials (0.3–3 ms post-stimulation) across distances aligned to peak response, at stimulation amplitudes of 5 µA, 25 µA, 50 µA, and 100 µA. (n = 25 probes across N = 9 recordings). **(g)** Quantification of the evoked potential magnitude measured at 1 ms post-stimulation as a function of distance from stimulation source and amplitude (n = 25 probes, N = 9 recordings, mean and SEM error clouds). **(h)** Quantification of evoked potential spatial extents superficial toward pia (negative values) and deep toward white matter and subcortical structures (positive values). Mean (large circle marker) and SEM (error clouds) spatial extents and individual data points (n = 25 probes). **(i)** Linear fits of the spatial towards the surface of the brain (dorsal, R² = 0.02) and deep to stimulation (ventral, R² = 0.16) **(j)** radial spatial extents for 50 µA stimulation for cathodal (cat.), anodal (an.), and bipolar (bi.) stimulation. Mean lines with SEM error clouds, and individual data points n = 50 (a ventral and dorsal spatial extent for each probe). See Figure S2 for full description of polarities and additional metrics.

Next, we empirically measured the spatial extent and magnitude of stimulation-evoked potentials using raw extracellular voltages recorded with Neuropixels probes (Figure 2e, Methods). We captured non-synaptic extracellular potentials that followed a brief, saturating electrical artifact and evaluated these in a window from 0.3-3 ms post-stimulation onset, a window that excludes the saturated portion (Figure 1l, Figure S1)

We characterized the extracellular potential for each probe (n = 25) across recordings (N = 9) superficial (or dorsal) to the stimulation source (denoted as negative distances) and deep (or ventral) to the stimulation source (denoted as positive distances). We aligned each probe to the peak response and averaged the evoked potential for 5, 25, 50, and 100 µA for cathodal (Figure 2f), anodal, and bipolar polarities (Figure S2); increasing stimulation amplitude led to larger evoked potentials, both in spatial spread and magnitude of potential, for all polarities (Methods). As expected, cathodal-leading monopolar (Figure 2f) and bipolar stimulation (Figure S2c) predominantly induced negative extracellular potentials, depolarizing membrane potentials [36, 46]. Anodal-leading monopolar stimulation leads to an early positive electrically evoked potential, consistent with a hyperpolarizing effect on membrane potential, but was followed by negative potential deflections beginning around 2 ms (Figure S2b). This observation aligns with previous literature suggesting early inhibitory effects of anodal stimulation but paradoxical somatic activations [36, 46–48].

To quantify the magnitude and spatial spread, we examined the extracellular potentials at a fixed time point of 1 ms, chosen to reflect a stable phase of the evoked potential across stimulation conditions (Figure 2g). We measured the magnitude of the evoked potential by calculating the mean maximum evoked potential and the area under the curve. For all polarities, both the maximum evoked potential and AUC increased nearly linearly (Figures S2g–S2h, see stats table). Next, we calculated the average spatial spread across all probes, both superficial and deep to the stimulation source (Methods). Both the superficial and deep distance from stimulation increased with amplitude (Figure 2h, superficial: 5 µA: -424.21 ± 209.65 µm, 25 µA: −683.18 ± 252.64 µm, 50 µA: −741.94 ± 267.22 µm, 100 µA: −743.84 ± 264.17 µm; Deep: 5 µA: 510 ± 214 µm, 25 µA: 591 ± 261 µm, 50 µA: 662 ± 263 µm, 100 µA: 712 ± 286 µm). However, contrary to predictions of the point source model in a uniform medium, the increase appeared sublinear and was poorly fit by linear regression (superficial R² = 0.02; deep R² = 0.17). Notably, boundary distances did not differ across polarities (Figure 2j, Figure S2e, 3-way linear ANOVA, F = 1.65, p = 0.20). This suggests the spatial limit of stimulation spread is primarily determined by amplitude rather than polarity, in contrast to some predictions suggesting more spatial focus to bipolar stimulation, but consistent with previous evidence and modeling expecting similar spatial extents [43, 46, 49]. All together, we found that the spatial extent increased sub-linearly with amplitude both superficially and deep from the stimulation source with no significant polarity effects.

To evaluate asymmetries in the spatial profile of evoked potentials, we compared the decay of the extracellular potentials measured (i) deep to the stimulation source towards a highly conductive but dense white matter boundary, and (ii) superficial to the stimulation source towards the saline bath on the surface of the brain (Figure 2h,i). The null hypothesis, provided by the isotropic point source model, would be no directional difference in the spatial extent measurements. However, the superficial and deep spatial extents were significantly different for cathodal stimulation (Figure 2i, 2-way linear ANOVA F = 1283.65, p < 0.001), and for all polarities (Figure S2e, 3-way linear ANOVA, F > 1000, p < 0.001, see stats table for full comparisons). This finding suggests that anatomical differences in boundaries, such as dense white matter or brain surface, affect the conductive properties of the tissue to create anisotropic spatial profile of the electrically evoked potential. We further explored this asymmetry relative to the underlying neural tissue using the alignment of evoked fields to mouse common brain coordinates.

### Anisotropic volumetric profile of the induced electrical field

To further explore the anisotropy and identify anatomical factors shaping the induced electric field (Figure 3a), we quantified the volumetric distribution of evoked potentials (Figure 3b, Methods). Volume, as measured as the 3D convex hull spanning the 1D extents recorded on orthogonal Neuropixels (Figure 2, Methods), trended to increase sublinearly with stimulation amplitude (Figure 3c; cathodal polarity: −25 µA: 0.31 ± 0.20 mm^3^, −50 µA: 0.43 ± 0.19 mm³, −100 µA: 0.61 ± 0.22 mm³; linear R^2^ = 0.259; see statistics table for full results across polarities).

**Figure 3.**
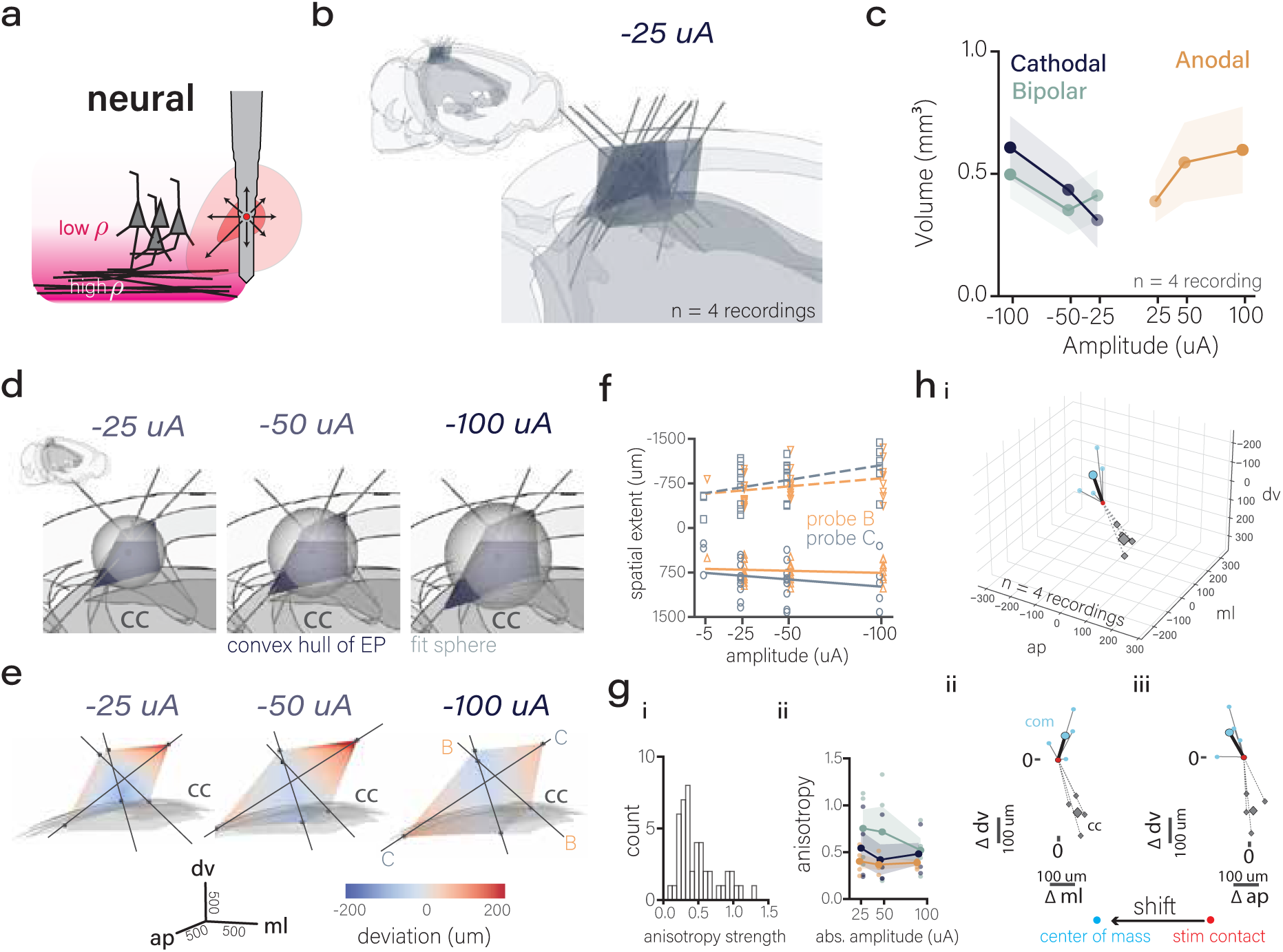
Anisotropic volumetric propagation of the induced electrical field. **(a)** Cartoon schematic depicting the alternative hypothesis, proposing that local differences in neural tissue resistivity and anatomical structures shape the evoked potential field. **(b)** 3D rendering showing volumetric extents of evoked potentials for -25 µA cathodal stimulation (n=4 recordings). gray mask represents corpus callosum and lines represent probe tracks. **(c)** Mean volume of evoked potentials across stimulation amplitudes (25 µA, 50 µA, and 100 µA) and polarities (cathodal, anodal, and monopolar) (n = 4 recordings, 2-way ANOVA, F(amplitude) = 0.16, p = 0.16, F(polarity) = 0.42, p = 0.16). Error bars represent standard errors of mean (SEM). **(d)** Example convex hull (purple) and spherical fit (gray) for one recording for -25 µA, -50 µA, and -100 µA. Inset of full brain provided for perspective. Gray mask (labeled CC) represents the corpus callosum **(e)** The mean deviation of the average convex hull of the evoked potential for 25, 50, and 100 µA cathodal stimulation compared to the isotropic convex hull derived from the fit sphere (n = 4 recordings, see Methods). Blue represents compressions from the null expectation and red represents expansions. Lines represent probe trajectories through the brain. B (orange) represents probe B which crosses through corpus callosum. C (blue) represents probeC where probe remains in cortex. **(f)** Upper (negative values, dorsal to stimulation, dashed line) and lower (positive values, ventral to stimulation, solid line) for probe B (crosses cc boundary) and Probe C (remains in cortex). R²(probeB dorsal) = 0.015, R²(probeB ventral) = 0.1, R²(probeC dorsal) = 0.05, R²(probeC ventral) = 0.187 (n = 9 recordings) anisotropy (see Methods for metric details) for all volumetric measurements across stimulus conditions. **(g)** anisotropy for each polarity (cathodal = blue, anodal = orange, and bipolar = green) by amplitude. Mean and SEM error clouds. **(h)** direction of anisotropy as measured by the direction and magnitude of the shift of the stimulation contact (red circle) to the center of mass (blue circles) of the evoked potential in relation to the closest point of the corpus callosum (cc, gray diamonds, see Methods) in 3D CCF space (i), and 2D slices (ii: ml-dv, iii: ap-dv). Larger markers represent means while smaller markers represent individual recordings (n = 4).

Inspection of reconstructed volumes in brain space revealed asymmetries at the gray–white matter boundary: evoked potentials extended broadly within cortex but were compressed ventrally at the corpus callosum (Figure 3d). To formally assess these asymmetries, we first established a symmetric reference by fitting the data with isotropic spheres (Figure 3d, gray spheres). This provided a baseline null expectation for isotropic evoked potential shape against which to compare the observed profiles. Because convex hull volumes underestimate extent along unsampled directions, we also constructed convex hulls from the fitted spheres, ensuring symmetry in both fit and measurement geometry. We then compared the observed convex hulls from evoked potentials to the imputed symmetric sphere-based hulls by calculating the Euclidean distance from each point on the symmetric hull to the nearest facet of the observed evoked potential hull (Figure 3e). This analysis extended the visual impression from individual examples (Figure 3d) to a condition-averaged measure of asymmetry. Deviations consistently localized to the ventral aspects of the fields: when probes remained in cortex (Probe C, Figure 3e), responses extended beyond the symmetric fit, but when probes traversed into subcortical white matter (Probe B, Figure 3e), responses were truncated relative to the sphere. To further quantify that observation, we directly compared the spatial extents of Probe B (which crosses the white matter tract) and Probe C (which remains in cortex) and found that there was a significant interaction between boundary (i.e., dorsal or ventral) and probe (multi-variate ANOVA, F(boundary:probe) = 6.46, p = 0.0123, see statistics table for full comparisons) confirming anisotropy that corresponds with surface and white matter boundaries.

We then used the volumetric fits from (Figure 3e) to quantify anisotropy. First, we computed the anisotropy strength (Figure 3f, Methods), defined as the norm of a directional bias term relative to the isotropic baseline radius. This dimensionless metric is 0 for perfectly spherical profiles and increases with directional variance. Across all amplitudes and polarities, anisotropy strength averaged 0.493 ± 0.288 and did not vary systematically with stimulation amplitude or polarity (Figure 3g, see statistics table). Second, we quantified the vector displacement between the stimulating contact and the center of mass (COM) of the evoked-potential volume relative to the corpus callosum (Figure 3h). The displacement magnitude exceeded zero (one-sample t-test vs. 0, p = 0.02), indicating anisotropy. The dorsal (superficial) component averaged 105 *±* 46 *µ*m away from the corpus callosum and, while directionally consistent, did not reach significance (one-sample t-test vs. 0, p = 0.1095). This superficial bias is consistent with ventral compression of evoked potentials at the white-matter (corpus callosum) boundary. Together, these analyses demonstrate that the induced electric field is not isotropic but shaped by anatomical constraints, including gray–white matter boundaries. Translationally, this anisotropy implies that in both DBS and cortical prostheses activation will preferentially spread along tissue planes and fiber pathways, and implies that current steering, electrode orientation, and patient-specific anatomy must be considered to achieve focal recruitment and stable percepts.

### Evoked spiking is sparse and increases in density with amplitude

Our measurements indicate that the potential produced by the induced electrical field from a stimulating electrode grows sublinearly with amplitude and is anisotropic, influenced by brain substrate. The neural responses of each neuron within this field, based on its morphology and physiology, may exacerbate or alleviate these spatial non-linearities and/or asymmetries. To understand the relationship between electrical stimulation and single neuron activation, we examined the spiking of single units directly following stimulation onset and how these responses relate spatially to the stimulation source, anatomical boundaries, and stimulation amplitude. Specifically, we measured single unit reliability, spatial uniformity and density, and across-class uniformity in response to electrical stimulation, and asked how the spatial distribution of activated units compared to the volume defined by evoked potentials.

To survey the evoked spiking, we first plotted each unit during the 10 ms following each −50 µA cathodal pulse, ordering units by cortical depth and annotating anatomical layers for each probe (Figure 4a, i–iii), and rasters from representative single units across the full amplitude range (Figure 4b, i–iii). Within this 10-ms window, we noted both spikes that occurred almost immediately after the pulse and spikes that were delayed by several milliseconds, the latter suggesting synaptically propagated activity. We sought to isolate spiking activity that was directly influenced by the electrical pulse and not derived from synaptic propagation, which we refer to as a “direct” response. We defined a direct response as spiking activity occurring within a 2-ms window immediately following stimulation onset. This window was chosen based on the distribution of first spike latencies, which exhibited a bimodal shape: an early peak immediately after stimulation, followed by a brief pause occurring at a median value of 1.95 ms across parameters and a broader, sustained peak beginning around 2.5 ms (Figure 4c, Figure S3). The 2-ms cutoff provides an estimate intended to capture most direct activation while minimizing spiking from any synaptically propagated activity. Historically, this early response has been difficult to measure due to saturating electrical artifacts and amplifier properties [50, 51]. However, in our data, we begin detecting spiking activity 200 µs following stimulation onset (Figure 4cii, Figure S3b), which closely aligns with the duration of signal saturation observed in our recordings (Figure 1l, Figure S1).

**Figure 4.**
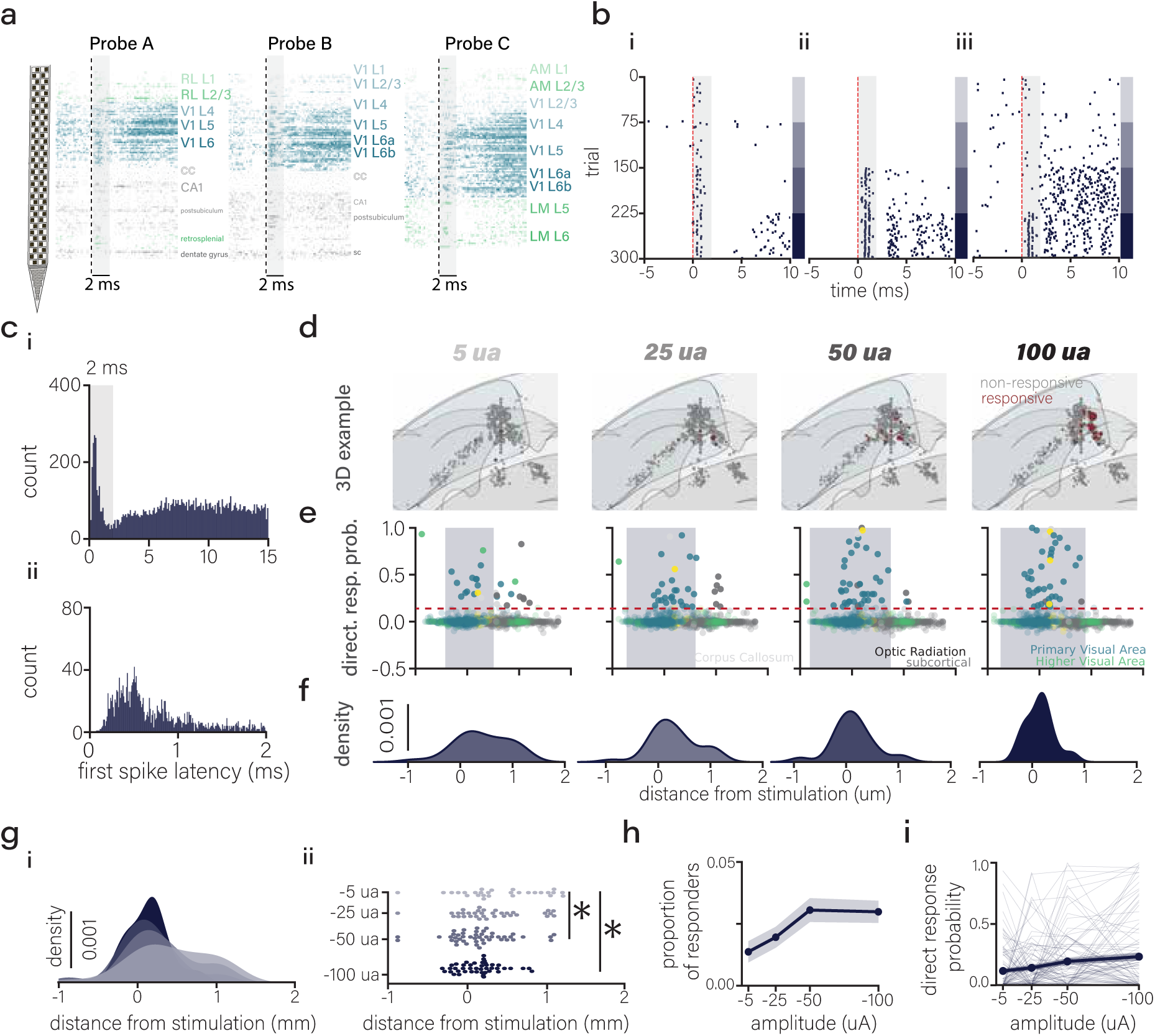
Evoked spiking is sparse and increases in density with amplitude. **(a)** Stacked raster for each probe trajectory for a 10-ms window with 2-ms direct response overlay (gray shaded) color coded by anatomical region sorted by depth for 50 µA cathodal stimulation. **(b)** Representative single unit rasters to increasing amplitudes (5, 25, 50, 100 µA) for a 10-ms window with 2-ms direct response overlay (gray shaded). **(c)** First spike latencies for 15-ms window with 0.1 ms bins (i) and 2-ms window with 0.01 ms bins (ii) for 50 µA cathodal stimulation (see Figure S3 for full polarity and amplitudes). **(d)** 3D renders of direct responders (red) and non-responders (gray) in CCF brain space for an example recording across 5, 25, 50, and 100 µA (blue shaded region represents V1). **(e)** Direct responder probability for single units across distance from stimulation, color-coded by anatomical brain region, across 5, 25, 50, and 100 µA for cathodal stimulation. **(f)** Direct responder densities across distance from stimulation for cathodal stimulation. **(g)** Densities (i) and distributions (ii) of direct responders across distance from stimulation for 5, 25, 50, and 100 µA. **(h)** Proportion of direct responders within the average evoked potential for each amplitude (n = 9 recordings, One-way ANOVA, p = 0.02). **(i)** Mean (thick line) and individual single unit (thin lines) direct response probability across amplitude for single units responsive at any amplitude (n = 99 single units, One-way ANOVA, p = 0.0004). See Figure S4 for direct response details for each polarity.

For each single unit, we calculated direct response probability, normalized for differences in firing rates (Methods). Single units with a direct response probability change following electrical stimulation greater than 0.15 were classified as “direct responders”. In visualizing responders in 3D anatomical brain space, such direct responders are sparse, amplitude-dependent, and while not spatially restricted, predominantly cluster within the cortex near the stimulation site (Figure 4d). To quantify the relationship between distance from stimulation and single unit direct response probability, we measured the Euclidean distance of each single unit from the stimulation source and measured the direct response probability as a function of distance from stimulation across amplitude (Figure 4e). These measures allowed us to address the density of neuronal activation within the applied field, and about the potential relationship between distance from stimulation and probability of direct activation.

Direct responders were sparse and increased in number (5 μA: 4.8±3.0, 25 μA: 5.7±3.5, 50 μA: 7.5±5.7, and 100 μA: 6.4±5.2 responders) and spatial density with amplitude (5 μA: 5.0±2.9, 25 μA: 13.2±11.1, 50 μA: 16.8±11.5, 100 μA: 11.9±5.8 responders/mm) (Figures 4e–4f). Direct responders were generally constrained within the average boundaries of the evoked potential (33.3% for 5 μA, 74.1% for 25 μA, 88.2% for 50 μA, and 100% for 100 μA, Figure 4e, shaded region). Notably, at the lowest amplitude (5 μA), more direct responders existed outside of the boundary compared to higher amplitudes, in line with previous observations [38] of antidromic or orthodromic somatic activation outside of the evoked potential from dendrites or axons near the stimulation source. Despite increases in the spatial extent of the measured evoked potentials (Figure 4e shaded region, from Figure 2), the spatial extent of single unit responders remained relatively constant across amplitudes and even decreased for 100 μA (5 μA: 1180±559 μm, 25 μA: 884±652 μm, 50 μA: 965±451 μm, 100 μA: 522±313 μm) (Figure 4g, ii). Stimulation amplitude significantly influenced the average distance from stimulation of responsive units (5 μA: 426 ± 500, 25 μA: 300 ± 436, 50 μA: 137 ± 394, 100 μA: 146 ± 260 μm, One-way ANOVA, F = 4.15, p = 0.007), specifically of 5 μA compared to 50 and 100 μA (Post-hoc Tukey tests, 5-50 μA p = 0.015, 5-100 μA p = 0.019, p > 0.05 for other pairs). Further, 5 and 100 μA recruited distinct spatial populations (pairwise two-sample Kolmogorov-Smirnov tests, corrected for six comparisons with Holm-Sidak procedure, 5 vs 25 μA: p_adj_=0.566, 5 vs 100 μA: p_adj_= 0.016, 25 vs 50 μA: p_adj_= 0.566, 25 vs 100 μA: p_adj_= 0.363, 50 vs 100 μA: p_adj_= 0.083, Figure 4g, i,ii). The proportion of responders within the evoked potential increased with amplitude (5 µA: 0.014 ± 0.12, 25 µA: 0.020 ± 0.14, 50 µA: 0.031 ± 0.172, and 100 µA: 0.030 ± 0.170, linear ANOVA: F= 5.41, p = 0.020, Figure 4h); and of the responders, direct response probability increased with amplitude (5 µA: 0.12 ± 0.20, 25 µA: 0.14 ± 0.21, 50 µA: 0.19 ± 0.25, and 100 µA: 0.23 ± 0.31, F = 12.78, p = 0.0004, Figure 4i), suggesting that increasing amplitude both recruits more cells and increases the spiking probability of recruited cells. All together, these results demonstrate that electrical stimulation induced direct neural firing in a sparse subpopulation of neurons, less than 5%, of the neurons within the spatial extent of the induced electrical field. While this population becomes denser with amplitude, the population of direct responders remains sparse. Next, we considered the composition of the sparse subpopulations directly activated by electrical stimulation near the stimulation site in cortex.

### Stimulation amplitude and polarity tune unit-specific recruitment

Because amplitude modulated not only how many neurons responded but where they were located, we next asked whether stimulation parameters (amplitude and polarity) could be used to selectively recruit single units. We expected that single units would exhibit a dose-dependent increase in direct response probability with increasing stimulation amplitude [36], such that the observed increase in density (Figure 4) resulted from the increased recruitment of a uniform underlying population. To test this, we clustered single units that were classified as direct responders at any amplitude, based on their direct response probability across amplitude (Figures 5a–5c, Methods), expecting most clusters to be dose-responsive; in fact, we have already shown on average in this subgroup of units, response probability increased with amplitude (Figure 4i). Hierarchical clustering produced 6 clusters (Figure S5a, cophenetic correlation coefficient (ccc) = 0.702, mean silhouette score = 0.352) that were broadly either dose-dependent or amplitude-dependent. Surprisingly, while 58.6% of direct responders exhibited dose-dependent increases in their response probability (Figures 5a–5c, i-iii, see stats table for stats on each cluster), we observed a subset of single units demonstrating amplitude selectivity, responding maximally, and almost exclusively, at specific intermediate amplitudes rather than monotonically increasing with amplitude (19.2% for 5 μA, 15.2% for 25 μA, and 7.1% for 100 μA, Figures 5a–5c, iv-vi, see stats table for stats on each cluster). We next asked whether stimulus polarity shapes which neurons are recruited. Clustering units by their normalized probability of firing to cathodal, anodal, and bipolar pulses (Figures 5d–5e, Figure S5b, Methods, n(clusters) = 6, ccc = 0.806, silhouette score = 0.519) revealed a striking polarity bias: 69.1% of directly activated units were polarity-specific, preferring bipolar (27.7%, Figure 5f, i), anodal (23.6%, Figure 5f, ii), or cathodal (17.7%, Figure 5f, iii) stimulation. The remainder fell into smaller groups that were largely insensitive to polarity: 10.9% were dose-selective but polarity-agnostic (Figure 5f, iv), 12.7% responded only when the cathodal phase of the pulse led (phase-selective, dose-dependent, Figure 5f, v), and 3.3% were driven exclusively by monopolar pulses (Figure 5f, vi; refer to the statistics table for a comprehensive breakdown of each cluster). These findings demonstrate that polarity and amplitude sharply tune single-unit recruitment, raising the possibility that intrinsic neuronal properties, beyond mere spatial location, affect which single units respond to particular electrical stimulation parameters.

**Figure 5.**
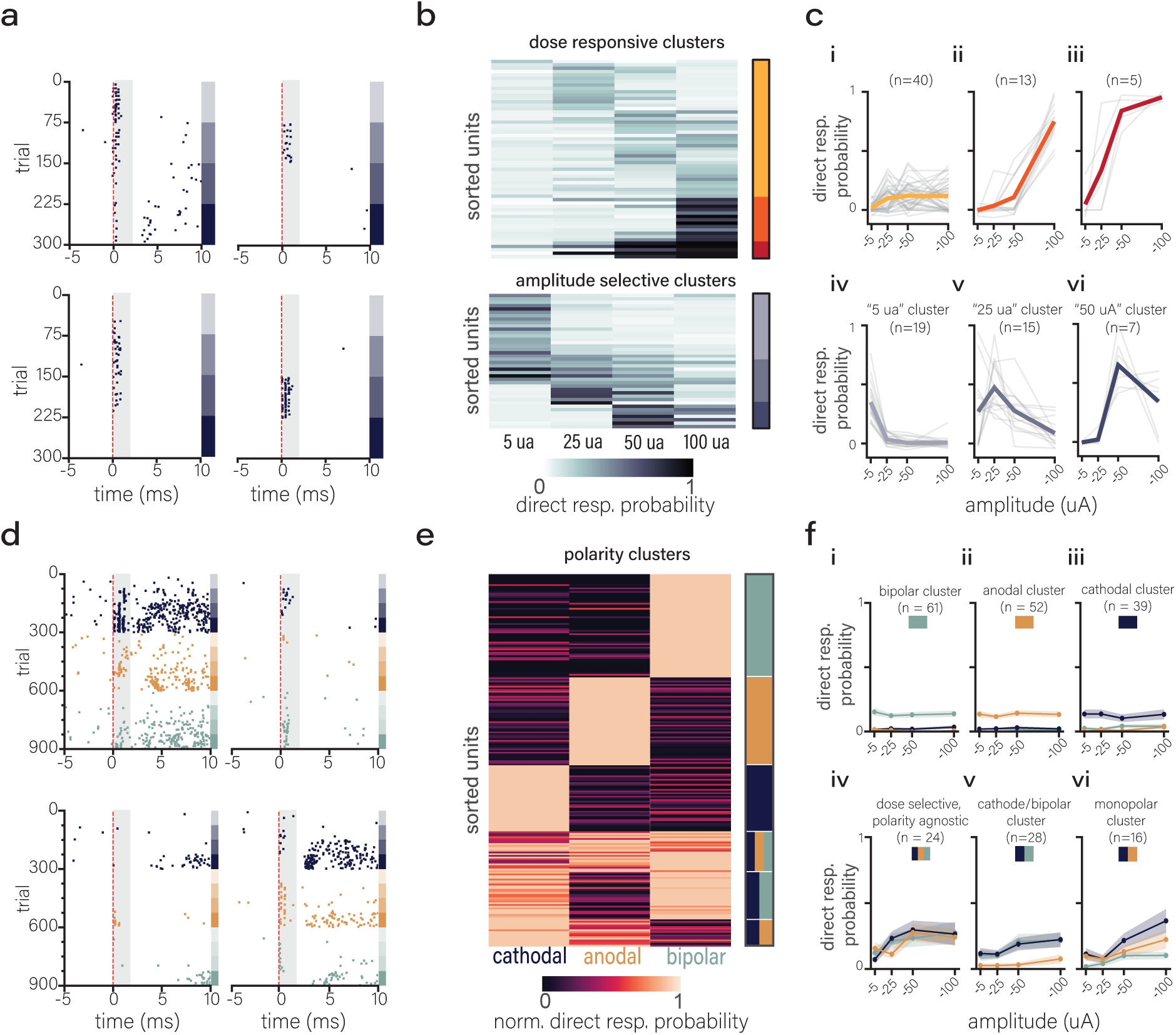
Stimulation amplitude and polarity tune unit-specific recruitment. **(a)** Rasters of example single-units showing select amplitude responsiveness (purple bars ordered by increasing amplitude). **(b)** Heatmaps sorted by labels from hierarchical agglomerative clustering of direct responders showing dose-responsive clusters (top) and amplitude-selective clusters (bottom) (cophenetic correlation coefficient (ccc) = 0.702, silhouette score = 0.352). **(c)** Direct response probability for single units of each cluster (i-vi) (color lines represent mean and gray lines represent individual single units). **(d)** Example single unit rasters showing select amplitude and polarity responses (bars on the right of rasters represent polarity and amplitude. Purple is cathodal; orange is anodal; and green is bipolar). **(e)** Heatmaps sorted by labels from hierarchical agglomerative clustering using polarity (ccc = 0.806, silhouette score = 0.519). **(f)** Direct response probability for single units of each cluster from the polarity clustering (i-vi). Mean lines and SEM error clouds. See Figure S5 for clustering statistics.

### Cell type influences direct activation distance distribution and response probability with amplitude dependence

To probe what makes some neurons, but not their neighbors, susceptible to specific pulses, we next compared how the two major cortical classes, fast-spiking (FS) interneurons and regular-spiking (RS) pyramidal cells, respond across space and stimulation parameters. We classified single neurons by the electrophysiologically recorded waveforms as FS, RS, or axonal [52] (Figures 6a–6b), and we analyzed the direct response of FS and RS cells. For RS cells, consistent with our findings across all single units (Figure 4e), increasing stimulation amplitude increased the number (5 μA: 15; 25 μA: 27; 50 μA: 34; 100 μA: 30 RS responders) and density (5 μA: 7.22, 25 μA: 13.74, 50 μA: 17.30, 100 μA: 26 responders/mm); however, for FS cells, increasing amplitude surprisingly did not increase the number of responders (5 μA: 12, 25 μA: 6, 50 μA: 10, 100 μA: 14) but did increase density by constraining the spatial distribution of those responders (5 μA: 11.23, 25 μA: 5.36, 50 μA: 28.55, 100 μA: 23.63 FS responders/mm) of FS responders (Figures 6c–6d). We observed differences in the spatial distribution of FS and RS responders at 5, 50, and 100 µA. In line with our previous findings (Figure 4), increasing amplitude constrained the average distance of stimulation for both FS responders (5 μA: 205 ± 311 μm, 25 μA: 203 ± 405 μm, 50 μA: -2 ± 122 μm, 100 μA: 60 ± 163 μm) and RS responders (5 μA: 567 ± 603 μm, 337 ± 455 μm, 208 ± 406 μm, and 185 ± 296 μm, 2 way ANOVA, F(amplitude) = 4.10, p = 0.008), but FS responders were clustered closer to the stimulation site than RS responders (2 way ANOVA, F(waveform shape) = 8.04, p = 0.005). FS responders were almost entirely constrained to the area within the evoked potential, even at low amplitudes (75.0% for 5 μA, 83.3% for 25 μA, 100% for 50 μA, and 100% for 100 μA of FS responders within evoked potential), while only 33% of RS responders were within the evoked potential at 5 μA, and that proportion increased with amplitude (74.1% for 25 μA, 88.2% for 50 μA, and 100 % for 100 μA, Figures 6c–6d). The differences among spatial distributions of FS and RS single units may reflect morphological and anatomical distinctions between the classes. These results point to the possibility that FS interneurons close to the stimulation site – with their compact axonal and dendritic arbors – are preferentially and locally recruited; while RS pyramidal neurons – with broader dendritic fields and longer-range projections – are more readily activated at a distance, revealing distinct spatial modes of excitability across cell types. To get a sense of how these spatial differences in the direct response (< 2ms) contributed to spiking which included early synaptic integration, we overlaid the spiking from all cortical units, color coded by waveform class and sorted by depth for a 10-ms window (Figure 6e) and averaged their firing rates (Figure 6f). Within this broader window, we noticed striking differences in spiking between waveform classes across amplitude which appeared to evolve (Kolmogorov–Smirnov tests at each amplitude: 5 μA: p_adj_ = 0.010, 25 μA: p_adj_ = 0.007, 50 μA: p_adj_ = 0.019, 100 μA: p_adj_ = 0.010, Figure 6f). These early differences raise questions about how the direct response from a single pulse of stimulation propagates through a network over longer timescales.

**Figure 6.**
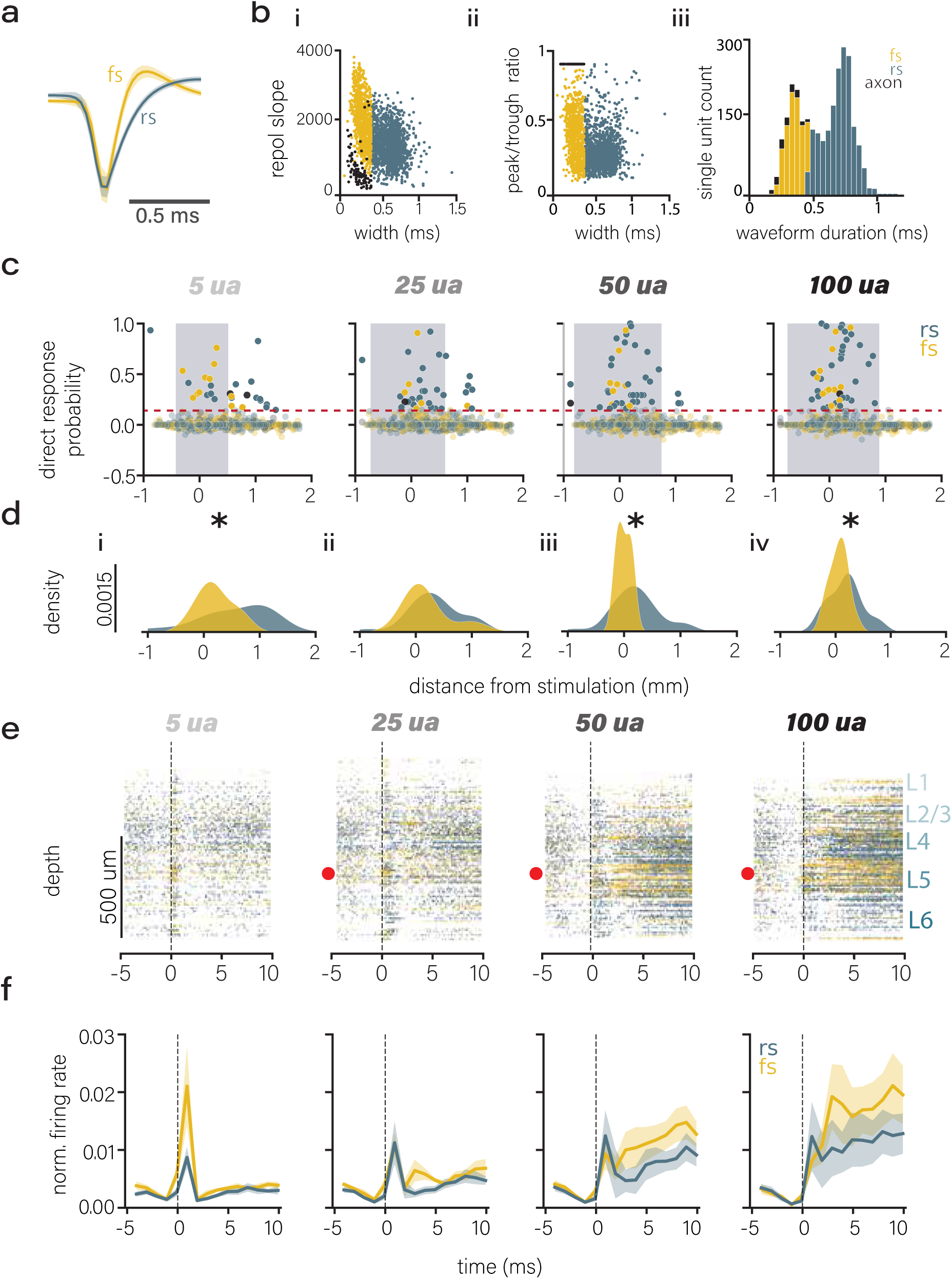
FS and RS cell type influences direct activation distance distribution and response probability with amplitude dependence. **(a)** FS (yellow) and RS (blue) waveforms (mean with SEM). **(b)** FS and RS waveform clustering based waveform duration (i-iii), repolarization slope (i), and peak to trough ratio (ii). **(c)** 5 µA: i, direct response probability for FS and RS single units across distance from stimulation (gray shaded region represents mean evoked potential) for 5, 25, 50, and 100 µA of cathodal single pulses. **(d)** responder density for FS and RS single units by distance from stimulation for each amplitude across 5, 25, 50, and 100 µA. **(e)** stacked rasters sorted by depth for FS and RS single units. **(f)** average normalized firing rates (bin width = 2 ms) for FS and RS single units (mean and SEM cloud, n = 9 recordings) across 5, 25, 50, and 100 µA. See Figure S6 for full polarity breakdown.

### Single pulse electrical stimulation leads to extensive circuit response

While the direct response to stimulation was sparse, with less than 5% of recorded neurons responding within the defined 2-ms window (Figure 4), this gave way to substantial amplitude- and region-dependent circuit activity. At higher amplitudes, stimulation triggered a global, highly synchronous response among cortical neurons that persisted for hundreds of milliseconds (Figures 7a–7b). This circuit-wide activity propagated asymmetrically, shaped by anatomical connectivity: subcortical units remained largely inactive, whereas neurons in visual cortical areas, well beyond the spatial reach of the initial stimulation, were robustly engaged (Figure 7b).

**Figure 7.**
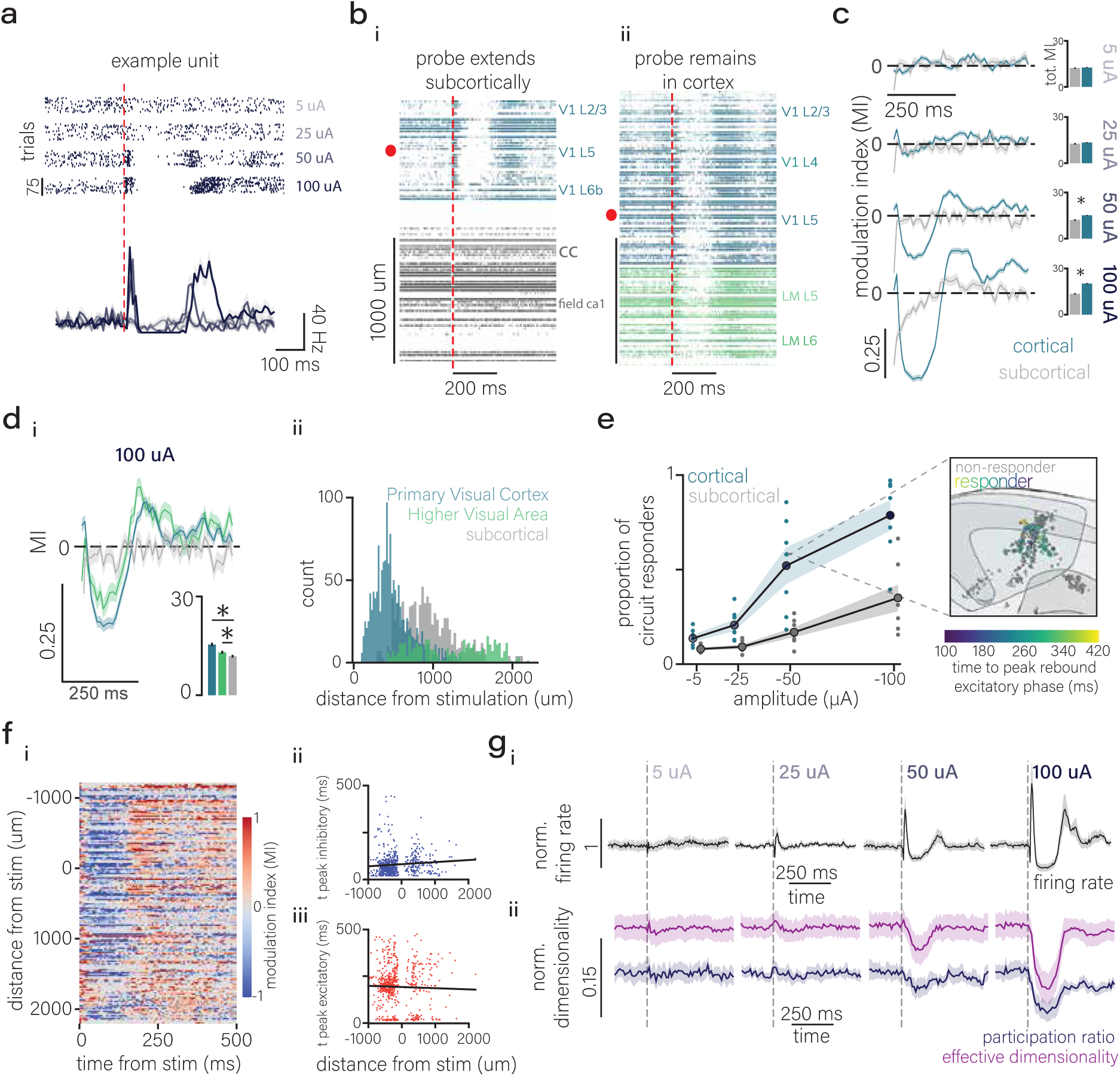
Single pulse electrical stimulation leads to widespread, asymmetric circuit responses. **(a)** raster (i) and averaged firing rate (ii) in response to single pulses of cathodal stimulation across 5, 25, 50, and 100 µA (mean and SEM overlay). **(b)** stacked rasters sorted by depths for example recording where probe extends to subcortical regions (i) and remains in cortex (ii) for similar insertion depths. **(c)** modulation indices (MIs) for a 500 ms window (5 ms bins) aligned to stimulation onset for cortical and subcortical single units across amplitudes for cathodal stimulation. Inset: cumulative modulation for cortical and subcortical (mean and SEM error). **(d)** i, MIs for primary Visual Area (V1), higher visual areas, and subcortical areas. ii, histograms of distances from stimulation for region area. **(e)** the proportion of responders for cortical and subcortical units for cathodal stimulation by amplitude. Inset: 3D render demonstrating circuit responders (color-coded by the time to their peak rebound excitatory phase) and non-responders (gray) from example recording for 50 µA cathodal stimulation. **(f)** i, Modulation Index heatmap for responsive cortical units sorted by distance from stimulation (negative values are dorsal to stimulation, positive values are ventral to stimulation). ii-iii, time to peak inhibitory response (ii) and time to peak excitatory response as a function of distance from stimulation **(g)** normalized firing rate (i, black), effective dimensionality (ii, purple) and participation ratio (ii, blue) across 5, 25, 50, and 100 µA (5 ms bins, mean and SEM clouds, n = 9 recordings). See Figure S7 and Figure S8 for additional circuit response analyses.

Cortical single units displayed a response consisting of early excitation, suppression, and at least one rebound excitation (Figure 7c), consistent with previous descriptions in the literature [19, 50, 53–55]. The magnitude of spiking modulation (Methods) increased with amplitude for cortical units (5 μA = 12.60 ± 8.43, 25 μA = 13.37 ± 8.41, 50 μA = 15.11 ± 8.80, 100 μA = 20.21 ± 10.32) but remained consistent for subcortical units (5 μA = 12.27 ± 7.21, 25 μA = 12.60 ± 8.00, 50 μA = 12.08 ± 7.53, 100 μA = 13.48 ± 7.70, Figure 7c, 2-way ANOVA: F(brain region) = 106.28, p(brain region)<0.001; F(amplitude)=487.36, p(amplitude)<0.001, F(amplitude:brain region)=90.88, p(amplitude:brain region)<0.001). The overall magnitude of modulation differed for cortical and subcortical units at 50 μA (Student’s T-test, p < 0.001) and 100 μA (Student’s T-test, p < 0.001, Figure 7c). This overall trend persisted for all polarities with minor differences (Figure S7d, multivariate ANOVA: F(polarity)=96.14, p(polarity)<0.001).

Notably, the differences in modulation was not solely driven by differences in distance from stimulation but rather connectivity; higher visual areas (HVAs) exhibited a response at higher amplitudes (magnitude of modulation = 15.82 ± 9.97) that was significantly different from sub-cortical units (magnitude of modulation = 13.48 ± 7.70) at 100 μA (Tukey-HSD, p = 0.004, see stats table for full comparisons, Figure 7d, Figure S8a) despite coming from a similar distance from stimulation distribution (Mann-Whitney, p=0.511, see stats table for full comparisons). This suggests that a single pulse of electrical stimulation in V1 is propagated along the cortical hierarchy. Further, the proportion of cortical neurons participating in this circuit response increases with amplitude (5 μA = 0.14 ± 0.05, 25 μA = 0.21 ± 0.08, 50 μA = 0.52 ± 0.26, 100 μA = 0.79 ± 0.21, F(amplitude) = 175.18, p(amplitude) < 0.001) such that at the highest amplitudes, over half of all recorded cortical units are recruited (Figure 7e). Despite being less responsive overall, the number of circuit responders in subcortical regions also increased with amplitude (5 μA = 0.08 ± 0.02, 25 μA = 0.09 ± 0.03, 50 μA = 0.17 ± 0.07, 100 μA = 0.35 ± 0.19, F(amplitude) = 175.18, p(amplitude) < 0.001; F(amplitude:region) = 15.61, p(amplitude:region) < 0.001; (F(brain region) = 40.14, p < 0.001) consistent with the volume increase we observed in the evoked potential (Figure 2), yet surprising given the spatial constraints of the direct response (Figure 4). There were small but significant differences across polarities, but overall they followed the same regional and amplitude trends (multivariate ANOVA, F(polarity)=5.19, p(polarity) = 0.006, see stats table for full comparisons, Figure S7c)

To address if this circuit response reflects a triggered, propagating wave, we sorted single units as a function of distance from stimulation and visualized their average modulation indices. Contrary to our expectation of a propagating wave, there was no apparent temporal structure, which was confirmed statistically by the lack of a relationship between distance from stimulation the time to peak inhibition (Figure 7f, ii, Pearson r = 0.09) or time to peak rebound excitation (Figure 7f, iii, Pearson r = -0.03). This suggests that these circuit dynamics likely reflect the synchronous recruitment of a recurrent circuit, either cortical [56] or corticothalamocortical [57, 58], that spans at least one millimeter, similar to the spatial scale of the evoked potential.

Finally, we sought to link circuit-wide responses to potential encoding capacity. To do so, we examined the dimensionality of neural population activity following electrical stimulation, because high population dimensionality is a hallmark of visual processing [59]. We explored population dimensionality using two Principal Component Analysis (PCA)-derived metrics: participation ratio, which reflects the evenness of variance distribution across principal components, and effective dimensionality which describes the number of dimensions required to explain 95% of variance (Methods). Despite the distinct temporal phases marked by fluctuations in firing rate (Figure 7g, i, black) that increased with amplitude, the response window was associated with a strong reduction in neural dimensionality, both participation ratio and effective dimensionality (Figure 7g, ii, purple and blue, Methods). This suggests that cortical responses to electrical stimulation, in contrast to responses to visual stimulation, become highly coordinated and individual neurons become stereotyped during the evoked response, even across periods of fluctuating firing rates. Notably, for 5, 25, and 50 μA, there were small increases in effective dimensionality within 20 ms of stimulation, which may reflect the window of optimal encoding before subsequent collapse of dimensionality. The impact of the dense increase in firing rate, concurrent with a collapse in dimensionality created diverging expectations for the behavioral accessibility of the neural activity driven by electrical stimulation.

### Single pulses of electrical stimulation are weakly detected, similar to low contrast visual stimuli

Given the extensive cortical effects of a single pulse of electrical stimulation, we sought to determine whether this extensive modulation of the cortex yields behaviorally relevant information without training for explicit detection of electrical stimulation. We first trained mice to detect visual stimuli. Then, we examined whether single pulses of electrical stimulation in V1 could be behaviorally reported like visual stimuli (Methods). Mice were trained to indicate detection of a stimulus by licking during a fixed response window (Figure 8a). Based on licking behavior, trials were classified as hits (rewarded detections), lapses (missed responses), or false alarms (licking before stimulus presentation; Figure 8b, Methods). Mice were initially trained to detect static gratings (Figure 8c, Phase 1), then we implemented a different visual stimuli, a “moving circle” stimuli to test the ability for detection to generalize (Figure 8c, Phase 2). Mice that successfully demonstrated detection of both visual stimuli classes moved onto Phase 3, where trials were intermixed, containing the previously trained visual stimuli as well as single pulses of electrical stimulation at varying amplitudes. This allowed us to directly compare the behavioral detectability of electrical stimulation pulses to visual stimuli of known contrast and perceptual characteristics (Figure 8c, Phase 3).

**Figure 8.**
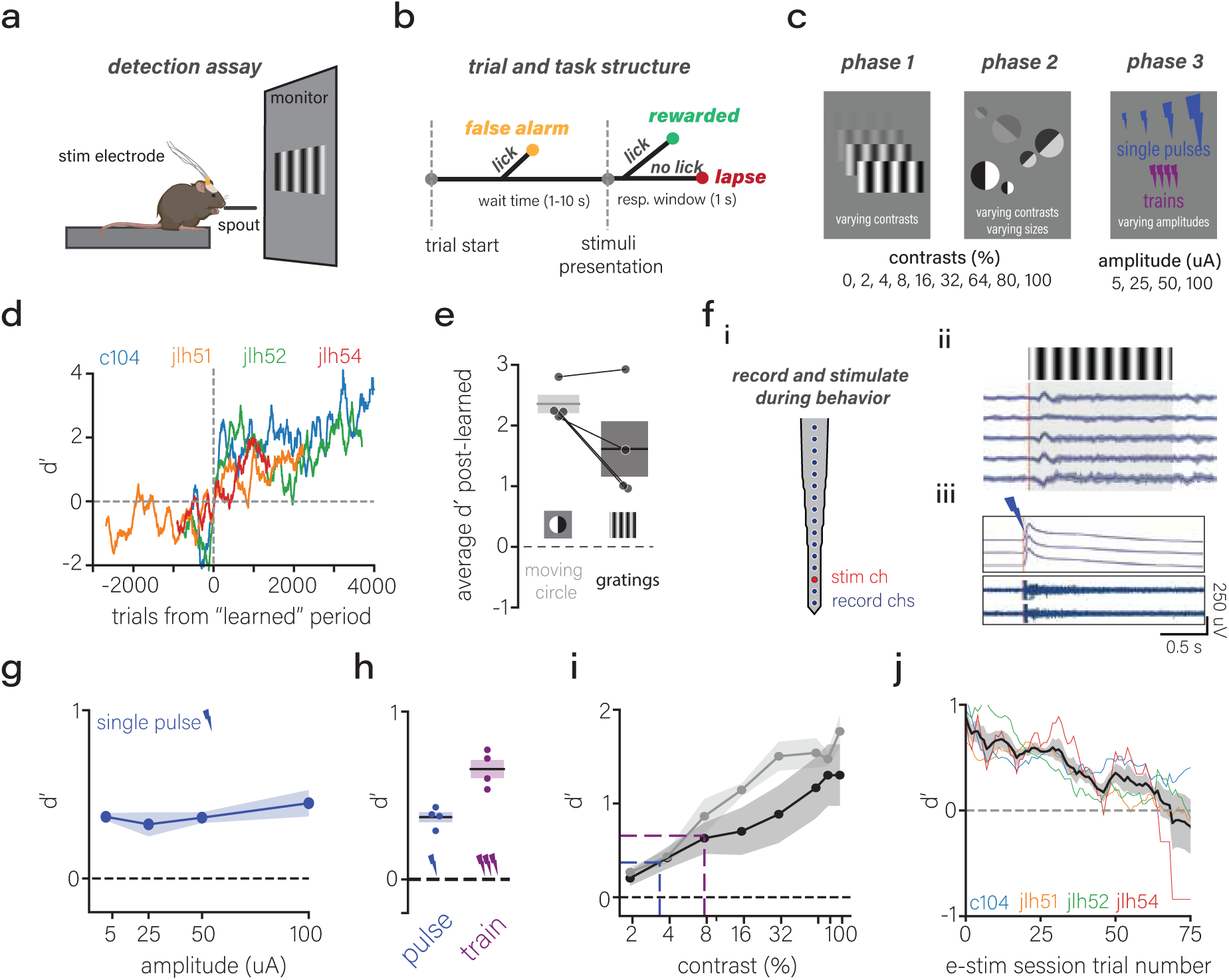
Single pulses are weakly detected similar to low contrast visual stimuli. **(a)** schematic of detection assay. **(b)** trial structure tree. **(c)** phase 1, static grating stimuli with varying contrasts. phase 2, addition of new visual stimulus, circles with varying contrasts and sizes. phase 3, addition of electrical stimuli with varying amplitudes for single pulses and trains **(d)** d’ for high contrast (>30%) gratings across trials aligned to the learned status. **(e)** d’ for high contrast static gratings and moving circle visual stimuli at the beginning of phase 3 (mean lines and SEM clouds. **(f)** i, schematic for recording and stimulating during behavior showing stimulation channels (red) and recorded channels (blue). ii, raw extracellular potentials from selected recorded channels in response to static gratings. iii, LFP (top) and AP (bottom) extracellular potentials showing response to a single pulse of electrical stimulation. **(g)** d’ across amplitudes of stimulation (cathodal and anodal pulses averaged, mean and SEM error clouds). **(h)** d’ for single pulses (blue) and trains (purple) (mean lines and SEM clouds). **(i)** d’ for static gratings (black) and moving circle (gray) across contrasts with mean line overlays for single pulses (blue) and trains (purple) (mean and SEM cloud overlay). **(j)** d’ across electrical stimulation trials within a session for individual mice (color-coded) and mean and SEM (black line). See Figure S9 for additional behavior analyses.

Mice successfully learned to detect high contrast static gratings (>30% contrast) over an average of 1353.8 ± 496.8 rewarded high contrast trials and 14802 ± 5154 total trials (n= 4 mice, Figure 8c, Methods). These mice generalized detection of static gratings to the “moving circle” stimuli before testing electrical stimuli, achieving a d’ of 2.35 ± 0.30, comparable to their performance on high-contrast grating stimuli (d’: 1.62 ± 0.92, p = 0.113). In one mouse, the moving circle task was introduced at a time when performance on high contrast grating stimuli exceeded a d′ of 6; this mouse immediately performed the moving circle task at a learned rate (d’ > 2), and within 500 trials, its d′ matched that of of the static grating stimuli (Figure S9b, i), suggesting the mice were capable of generalizing to different visual stimuli without explicit training.

Following visual training, mice were chronically implanted with stimulating electrodes (Neuronexus, CM16) in V1, using the same stereotactic coordinates and depths validated in the previous acute electrophysiological recordings. To confirm that electrodes were accurately positioned within visually responsive cortical areas and functioning, we simultaneously recorded from non-stimulating contacts on the implanted stimulating electrode during behavioral sessions (Figure 8f, i). Recorded channels exhibited visually evoked potentials to high-contrast visual stimuli (Figure 8f, ii), verifying placement within visually responsive cortical regions. Additionally, we verified effective delivery of electrical stimulation by recording neural activity from non-stimulated electrode contacts during stimulation trials. As expected, we observed large electrical artifacts coinciding precisely with stimulation events in both LFP (Figure 8f, iii, top) and AP bands (Figure 8f, iii, bottom). On specific contacts, we noted increases in multi-unit activity following stimulation, confirming electrical pulses reliably modulated neural activity during the task (Figure 8f, iii, bottom).

Mice detected single pulses of electrical stimulation at similar rates for all amplitudes (d’ for 5 μA: 0.37 ± 0.04, 25 μA: 0.32 ± 0.14, 50 μA: 0.36 ± 0.06, 100 μA: 0.45 ± 0.14; F(amplitude) = 1.74, p = 0.21, Figure 8g), and above the level of chance (average pulse d’ = 0.37 ± 0.06, one-sample T test compared to d’ of 0, p = 0.001, Figure 8h). Anodal leading waveforms were marginally more detectable than cathodal leading (anodal d’ = 0.44 ± 0.06, cathodal d’ = 0.33 ± 0.09, paired T-test, p = 0.028, Figure S9c). High-frequency stimulation trains (100–300 Hz; 10–50 pulses), similar to those commonly used in visual cortical prostheses [20], yielded higher detection sensitivity than single pulses (d′ = 0.66 ± 0.11 vs 0.37 ± 0.06; paired t-test p = 0.019). Within trains, detection did not differ by frequency (100 Hz 0.72 ± 0.13; 200 Hz 0.46 ± 0.14; 300 Hz 0.61 ± 0.09; one-way ANOVA F = 3.22, p = 0.112), though frequencies were unevenly sampled. Performance on visual stimuli for static gratings (2%: 0.20 ± 0.16, 4%: 0.42 ± 0.23, 8%: 0.63 ± 0.33, 16%: 0.70 ± 0.50, 32%: 0.89 ± 0.61, 64%: 1.17 ± 0.65, 80%: 1.30 ± 0.66, 100%: 1.30 ± 0.65) and moving circles (2%: 0.27 ± 0.11, 4%: 0.43 ± 0.14, 8%: 0.86 ± 0.37, 16%: 1.15 ± 0.14, 32%: 1.50 ± 0.31, 64%: 1.54 ± 0.31, 80%: 1.48 ± 0.15, 100%: 1.77 ± 0.35) increased with contrast (static gratings: One-way ANOVA, F(contrast) = 18.28, p < 0.001; moving circle: One-way ANOVA, F(contrast) = 46.52, p < 0.001; Figure 8i). Perceptual detection in the highly trained mice began at 4% contrast for static gratings (one sample T-test: p = 0.035) and 2% contrast for moving circles (one sample T-test: p = 0.018, Figure 8i). Single pulses of electrical stimulation performed similarly to a visual contrast between 2 and 4% while trains performed near 8% contrast (Figure 8i). Interestingly, within a session, electrical stimulation became less detectable across trials (Mixed Linear Model regression: y-intercept = 0.56, p(intercept) < 0.001, z(intercept) = 6.61; slope = -0.004, p(slope) < 0.001, z(slope) = -15.18, Figure 8j), suggesting the mice became habituated or desensitized to repeated presentations of electrical stimulation, consistent with previous reports in both somatosensory and visual cortex [21, 60]. Reaction times decreased with increases in contrast for both classes of visual stimuli, while electrical stimuli reaction times were similar to those of low contrast visual stimuli (Figure S9f, see statistics table), suggesting electrical stimuli are low salience. Lastly, it’s a common phenomenon that percepts evoked by electrical stimulation are unnatural but can be learned over time [61]. Our behavioral paradigm was not designed to test learning of electrical stimulation, but rather to assess its innate detectability and generalization to visual stimuli. Nevertheless, because electrical stimulation was tested across many sessions, the mice had ample opportunity to improve with experience. We found no evidence of such improvement: detection performance did not increase across sessions (Ordinary Least Squares, R² = 0.01, F = 1.12, slope = –0.004, p(slope) = 0.29, intercept = 0.39, p(intercept) < 0.001).

Overall, we found that mice weakly detected electrical stimulation, similar to that of low contrast visual stimuli. These findings suggest that single pulses of electrical stimulation, despite inducing widespread cortical modulation persisting hundreds of milliseconds, are only weakly detectable perceptually: a disconnect between cortical activation magnitude and behavioral salience.

## Discussion

Electrical stimulation is a cornerstone of both basic neuroscience [1, 47, 62, 63] and clinical neuromodulation [4, 5, 9, 22, 23, 25, 64, 65], and significant effort has been made to understand how electrical stimulation interacts with neural tissue [36, 37, 48, 66–68]. Despite widespread use, several central questions remain regarding the mechanisms and effects of electrical stimulation, including but not limited to the volume of neural tissue activated, spatial symmetry, cell-type specificity, and functional impact.

We measured the volume of neural tissue, and the individual neurons, that are affected by single pulse electrical stimulation. Classic work has argued that the volume of activated tissue expands with stimulation amplitude, inferred from measuring the spiking activity of a single pyramidal neuron to varying amplitudes[37]. Computational modeling of electric fields, largely in uniform media, has supported this view, indicating that the size of the electric field increases with the amount of current injected [39, 68] (and see Figure 2b). Congruent with this, optical measurements using voltage dyes [69] report millimeter-scale changes from single pulses of electrical stimulation. In contrast, using 2-photon calcium imaging to infer neural spiking in cortex, Histed [38] inferred that directly driven spikes arise from a compact zone whose size is relatively insensitive to amplitude, with higher currents primarily increasing the density of recruited neurons, not the volume. The somatic locations of the activated units are sparsely dispersed, activated from axons and dendrites within tens of microns from the stimulation source. To reconcile the apparently conflicting conclusions [37, 38], investigators have turned to anatomically detailed, biophysically grounded intracortical microstimulation models [39]. This modeling found the effective field expands with amplitude, whereas the somatic activation volume is amplitude-invariant and fills in with additional units. Here, we used Neuropixels to measure both the spatial extent of the applied electric field and the somatic activation. Consistent with voltage-sensitive dye imaging [69], evoked potentials extended 0.5 to 1 mm in radius and scaled sub-linearly with amplitude, matching predictions from modeling [68] and several lines of experiments measuring direct activation of cortex [37, 56]. Yet within this evoked potential, directly activated single units did not expand with current; instead, higher amplitudes increased the density of responsive neurons within a fixed space, closely matching the “constant-radius/increasing-density” picture [38]. Together, these results support the proposal that amplitude-dependent electric field can coexist with sparse, spatially constrained neural spiking [39].

Previous experimental attempts to measure electrically-evoked neural activation have been limited by stimulation artifacts that obscure the earliest responses, by limited spatial context during electrophysiology, and/or by the temporal resolution of imaging techniques [70]. Neuropixels probes help overcome limitations from electrical artifacts through band-specific amplifier gain settings[40] and, we found, integrated amplifiers with rapid recovery that allow observation of neural activity within 300 µs after pulses even at high amplitudes and within hundreds of microns from the stimulation source. The orthogonal geometry in our experiments and alignment to Allen CCF brain space [41] allowed three-dimensional sampling, overcoming historical difficulties with spatial information with electrophysiology. This combination – millisecond-scale timing, high density sampling, and 3D spatial registration – provides a framework to connect evoked potentials, direct single unit activation, and circuit-wide activity.

Computational models of electrical stimulation typically assume a uniform excitable medium even when incorporating cytoarchitecture [39, 42, 49, 66, 68]. However neural tissue is structurally and biophysically heterogeneous: conductivities differ across gray vs. white matter and across layers [44, 45]. Further, axons, especially at the axon initial segment, reach threshold more readily than somata [36, 46, 71] and myelination and ion-channel composition shift excitability both within a cellular compartments [66, 72–74] and across cell classes [75, 76]. Finally, morphology/orientation govern field–membrane coupling [36, 43, 75, 77]. While thresholds for specific elements can be measured in isolation, extending those measurements to population-scale *in vivo*, where diverse elements and orientations intermix, is not tractable. Our data show that anatomical heterogeneity, both broadly from influences from white-gray matter boundaries (Figure 3) and from cellular diversity (which may encompass morphological and diversity in ion channel expression)(Figure 6), creates anisotropic neural responses to electrical stimulation. White-matter boundaries limited evoked potential propagation. At a cellular level, fast-spiking (putative interneuron) and regular-spiking (putative pyramidal) units exhibited distinct spatial patterns. These differences may arise from their distinct morphologies [78–80], but also may be influenced by morphoelectric properties derived from differences in ion-channels and conductances [75, 76, 81, 82]. Additionally, increasing amplitude increased recruitment density rose and the spatial profiles of fast- and regular-spiking populations partially converged, consistent with a shift from predominantly axono-/dendritic activation to somatic activation. We also observed polarity- and amplitude-selective units, suggesting that complex interactions that occur in heterogeneous brain tissue, including those between cellular morphology, electrophysiological properties (e.g., ion-channel distributions, membrane resistivity), and orientations to stimulation, create amplitude- and polarity-selective responses in some neurons beyond simple dose-dependence that may be expected [83, 84]. This specificity has strong implications for the application of electrical neuromodulation – that the parameters of such modulation could be algorithmically controlled to enhance the activity of certain neurons over others by adjusting polarity and amplitude, resulting not only in changing the number of neurons in stimulated ensembles (Figure 4d), and their level of activation (Figure 4b), but also the constituents of those ensembles (Figure 5b). Future work should take a principled approach to discovering parameters that can drive meaningful differences in neural ensemble activation. Collectively, these findings highlight the role of anatomical and cellular heterogeneity in determining neural responses to electrical stimulation, which could be exploited in future efforts to precisely control information delivery as in sensory prosthetics.

Electrical stimulation has been successfully used to elicit percepts for decades, from somatosensory tingles [13, 85] to visual phosphenes [23, 86], yet the resulting sensations remain coarse and unnatural [29]. Most strategies to improve perceptual outcomes scale electrode count and/or coverage across representational space in early sensory cortices [87–90]. However, in V1 and other cortical areas multiple maps of features are overlaid, with areas and even single cells encoding multiple features and mixed information [91, 92] and perception depends on distributed, time-evolving population activity [33, 93]. Our results show that despite eliciting a large, synchronous cortical response that spreads across V1 and higher visual areas, and persists for hundreds of milliseconds, single-pulse microstimulation was weakly detected across all amplitudes. This dissociation implies that the strength and number of neurons activated are not limiting factors for sensory perception. Instead, perception of artificial inputs may require matching distributed, temporally-structured patterns of activation.

Visual stimuli recruit distributed, temporally structured patterns in primate [93–95] (Gallant fMRI, Murray/Wang 2012) and mouse V1 [96], whereas single pulse electrical stimulation elicits a highly synchronous wave of activity and reduces population dimensionality, despite fluctuations in firing rates(Figure 7). This neural difference is consistent with perceptual observations: reliable detection of electrical stimulation in V1 typically requires extensive training and trains of many electrical pulses [20, 97–100]. Such detectability may reflect a learned associations rather than native neural interpretability of evoked activity, as demonstrated down to single neuron activation detection [101, 102]. In our study, we found that without explicit training, artificial stimulation of V1 is not innately behaviorally salient: mice successfully generalized rapidly to novel visual stimuli but did not generalize to electrical stimulation. Relatedly, work in non-human primates has demonstrated that training to detect V1 microstimulation impaired visual contrast detection, and conversely, retraining on visual contrast impaired microstimulation detection [100]. Because visual and electrical stimulation trials were intermixed during our testing phase, this interference may have limited perceptual learning of electrical stimulation in our paradigm. Further, in sensory cortical prostheses, humans often report habituation to percepts, where percepts become less detectable over repeated deliveries [21]. We noted habituation to detection of electrical stimuli within a behavior session, despite no apparent changes in the evoked neural activity across stimulation pulses, suggesting that the stimuli lose perceptual salience, potentially through reductions in adaptation to evoked V1 outputs. Moreover, when electrical stimulation is used to bias ongoing sensation (e.g., MT microstimulation shifting motion judgments [1], effects are robust even with small currents, reinforcing perceptual salience depends more on naturalistic cortical dynamics than the absolute magnitude of driven activity [103].

In this study, we chose to limit measures to the neural responses to single pulses of electrical stimulation, despite widespread clinical and experimental applications employing trains of pulses (typically 100–300 Hz, 30–100 pulses). Our design isolates the neural effects of individual stimulation events without the confounds of temporal summation of high-frequency stimulation [84]. By characterizing the fundamental unit of electrical stimulation – single pulses – we provide a set of bases for predicting the neural effects of more complex stimulation paradigms from trains to biomimetic paradigms that use spatiotemporally patterned pulses [22].

Our findings motivate several avenues for improvement of neural devices or approaches for augmentation and addition of neural information to cortical circuits. Directly comparing the statistics of naturalistic neural dynamics to artificial stimuli may provide quantitative benchmarks for prosthetic design that inform spatiotemporally-patterned modulation. Exploring alternative stimulation waveforms (beyond biphasic pulses, such as sinusoidal or other continuous forms) may further unlock regimes of sensory neuromodulation. Future work could leverage machine learning approaches to optimize stimulation parameters for neural or behavioral targets—for example, reproducing cortical response patterns of natural visual stimuli or enhancing detectability in behavioral tasks – a strategy that is beginning to be explored [104]. Further, stimulation using multiple electrode contacts – arranged not only horizontally but also across cortical layers, and delivered with temporal patterns – may be essential for recreating high-dimensional visual inputs [59, 105] that reflect both the temporal dynamics of neural spiking and anatomical connectivity. Further, stimulation from multiple contacts, not only across horizontal brain space but laminar space, and temporally varied may be essential to recreating high-dimensional visual inputs with consideration to the temporal flow of spiking and anatomical connectivity.

Together, our results suggest the amplitude of neural activation does not straightforwardly predict perception. Instead, the spatiotemporal pattern of activity is a determinant of perceptual impact. For cortical prosthetics and other neuromodulatory technologies, this implies that more current or more synchronous firing is not necessarily better; rather, effective stimulation will require tailoring inputs to mimic the spatial and temporal structure of natural neural signals. Advances in technology (e.g., high-density electrode arrays, current steering, adaptive closed-loop control) and leveraging anatomical heterogeneity for selective neural responses open the door to such biomimetic stimulation strategies. Ultimately, understanding the principles of how electrical stimulation engages neural circuits and leveraging these principles in neuromodulatory devices will be essential for building neuroprostheses capable of delivering reliable naturalistic percepts.

## METHODS

### Surgical methods

Surgical procedures included headbar affixation and habituation, craniotomies, and chronic implantation of linear 16-channel electrode arrays.

#### Headbar affixation procedure

Mice underwent surgery to fix a metal headbar to the skull for head fixation. Mice were anesthetized with 5 % isoflurane and placed on a stereotactic frame (Model 940, Kopf Instruments). Isoflurane was maintained at 1.5-2% (Somnosuite, Vetflo) and body temperature was held at 37^◦^C using a temperature controller (TC-1000, CWE). Anesthesia depth and mouse wellness were checked every 15 minutes, ensuring a stable breathing rate between 15-20 breaths per minute, temperature, and the absence of a toe pinch reflex. Carprofen (5 mg/kg) and lactated ringers were delivered for pain management and surgery recovery. The hair above the skull was removed, and the surgery site was sterilized with iodine and ethanol scrub. A portion of the skin and periosteum above the skull were removed. The skin surrounding the skull was secured with VetBond adhesive glue. The skull was cleaned and dried with phosphate-buffered saline. The skull was leveled by aligning lambda and bregma in the anteroposterior and mediolateral axes. The headbar was placed on the skull with the headbar opening approximately centered on the skull above the left primary visual cortex. The headbar was fixed to the skull using translucent CI&B Metabond (Parkell) dental cement. Once the dental cement dried, mice were returned to a home cage with a heating pad and moist chow and monitored for 30 minutes or until full recovery.

#### Head-fixation habituation

Following headbar affixation, mice were allowed at least one week to fully recover. Mice were habituated to head fixation for acute electrophysiological recordings and behavior (see behavior section for behavior-specific habituation). For the first day, mice were handled and allowed to explore the head-fixation apparatus freely. On subsequent days, mice were head-fixed starting with 15 minutes and progressing to 90-120 minutes. Habituation sessions were terminated if mice exhibited distress, including vocalizations, physical agitation, and porphyrin secretion in the eye. The recording conditions were simulated during later habituation sessions, including exposure to direct light from the stereoscope and the high-frequency noise of the piezoelectric micromanipulators (New Scale Technologies). Following each habituation session, the mice were given peanut butter as a reward.

#### Craniotomy for acute electrophysiological recording and chronic Neuronexus implantation

Craniotomies for acute electrophysiological recordings or chronic multichannel stimulating electrode (see “Visual and electrical stimulation behavioral detection assay” subsection) implantation were performed on mice that previously underwent the headbar affixation procedure. Mice were anesthetized and monitored as described in the headbar affixation procedure. Lambda and bregma were identified through the clear dental cement by applying a thin layer of saline over the dental cement. The skull was leveled, and stereotactic coordinates were identified. For acute electrophysiological recordings, a small craniotomy (2x2 mm) encompassing each insertion stereotactic coordinates or individual burr holes at the predetermined stereotactic coordinates for each probe were performed. In each case, the dura was removed to facilitate probe insertion. For chronic multichannel stimulation electrode implantation (Neuronexus A1x16-5mm-50-703-A16, plated with IrOx for more effective current delivery), a small burr hole with dura removal at (-3860 AP, -2580 ML) was performed. The multichannel stimulation electrode was held using a custom 3D printed part and inserted 900 μm into the brain. The burr hole and exposed shank were surrounded with Kwik-Sil (World Precision Instruments), and the electrode base was secured to the headbar with clear CI&B Metabond (Parkell) dental cement.

#### Stereotactic coordinates

| Electrode | Entry (AP, ML, DV) | Tip (AP, ML, DV) | Angles | Depth ($\mu\text{m}$) |
| --- | --- | --- | --- | --- |
| Stim | (−3407, −2504, −202) | (−3407, −2504, −1602) | (0, 90, 0) | 1200 |
| Probe A | (−2839, −2764, −276) | (−4017, −1360, −2109) | (130, 45, 0) | 3000 |
| Probe B | (−3514, −2310, −135) | (−1709, −2310, −1939) | (0, 45, 0) | 3000 |
| Probe C | (−2990, −2271, −104) | (−4040, −4166, −2271) | (−119, 45, 0) | 3000 |

### Electrophysiological recordings

Electrophysiological recordings used multi-Neuropixels with concurrent electrical stimulation exploring a vast parameter space.

#### Electrophysiology data collection

Neural recordings used Neuropixels 1.0 probes[40] with the 383 electrode sites closest to the tip active. Signal for each recording site (channel) is split into a spike band (30 kHz sampling rate with a 500 Hz first-order highpass filter) and an LFP band (2.5 kHz sampling rate, 1000 Hz lowpass filter). The experimental rig was assembled using Thorlabs pieces and designed to insert 3 Neuropixels at 45-degree angles orthogonal to a stimulating electrode inserted at 90 degrees. Neuropixel probes and stimulating electrodes were mounted on 3-axis piezoelectric micromanipulators (New Scale Technologies). The New Scale manipulators were secured to a round breadboard attached to a 2D microscope platform (Thorlabs) to allow fine X and Y motor control. Probe geometry and relative spacing between probes were set before insertion and retracted at least 1000 μm above their target insertion depth. The mouse platform with an adjustable height was situated under the probe platform and raised to the appropriate height for insertion. X and Y positions were adjusted using the X,Y microscope platform to center probes above the insertion points. Probes were lowered to brain depth and then inserted to target depths (between 2000-3000 μm for Neuropixels and 1000 μm for stimulating electrode) at a rate of 50-100 μm/min. Neuropixel data was acquired using Open Ephys GUI software[106].

#### Electrical stimulation hardware and software

Electrical stimulation was delivered using a combined waveform generator and isolated current output device (AM4100, A-M Systems). Stimulation during electrophysiological recordings was controlled with a modified version of example Matlab code from AM-systems. This code used serial commands to update parameters, trigger stimulation, and save stimulation parameters as CSVs. Stimulation for the behavioral detection task was controlled using custom Python code interfacing with the stimulator using PySerial commands. Matlab and python code for stimulator interface is available (github.com/denmanlab/am4100_code).

Stimulation was delivered through contacts on a linear 16-channel Neuronexus electrode (Neuronexus, A1x16-5mm-50-703). To increase the stimulation efficacy, each site was activated with Iridium Oxide. The contact size is 703 μm^2^m with 50 μm spacing between contacts. As previously noted, the average contact impedance was 0.10 *±* 0.06 M (Figure 1j, top), and the average contact charge capacity was 25.90 *±* 9.69 mC/cm^2^ (Figure 1j, bottom).

#### Electrical stimulation parameters

Stimulation parameter space was systematically explored. Each unique stimulation parameter was delivered 75 times with at least 2 seconds between single pulses or trains. Each pulse was a biphasic square wave, where the sign of the amplitude indicates the phase of the first part of the wave. Each recording tested a range of amplitudes including -100, -50, -25, -5, 5, 25, 50, and 100 μA and monopolar and bipolar polarities. Negative amplitudes indicate cathodal-leading pulses and positive amplitudes indicate anodal-leading pulses. “Bipolar” stimulation is used to describe cathodal-leading bipolar configurations; “anodal” and “cathodal” is used to describe anodal-leading and cathodal-leading monopolar configurations, respectively. For the analyses in the text, we tested single pulses with a pulse width of 100 μs for each phase from contacts in or near Layer 5 of primary visual cortex. Additional stimulation parameters were tested in select recordings—including varying frequencies, stimulation depths, and pulse widths—that are not included in the analyses but are available in the dataset. In the text, we report the current magnitude in amperes, but provide conversion to charge density (C/cm²) for our electrode configuration to help extrapolate our findings (Figure 1k). We used the following calculation for charge density:

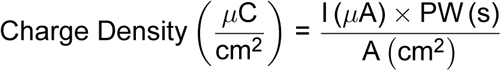

#### Spike sorting

Raw spike-band data was input into Kilosort 2.5 (https://github.com/MouseLand/Kilosort), an automated template-based spiking sorting algorithm[107]. Following spike sorting with Kilosort, the clustering and classification of single units were manually curated using Phy (https://github.com/cortex-lab/phy). Additional quality metrics were performed to ensure quality of clustered single units and are described in the “Quantification and statistical analysis” section.

### Visual and electrical stimulation behavioral detection assay

We trained mice to report detection of visual stimuli with different classes and then tested to see if mice that were proficient at reporting detection of visual stimuli could also detect electrical stimulation of primary visual cortex. Here we describe the task protocol; trial structure; task design and stimuli (visual and electrical); task software and hardware; and the approach for electrophysiology during behavior sessions. The statistical analyses will be described in the subsection “Quantification and statistical analyses”.

#### Task protocol

Mice were surgically implanted with a headbar for head-fixation and allowed one week of recovery before behavioral training (see “Head affixation procedure” subsection). Following recovery, mice were weighed daily for three consecutive days to establish a stable baseline weight. Mice were placed on a controlled water deprivation schedule, receiving approximately 1 mL of water daily to maintain their body weight at approximately 85% of their baseline, adjusted for expected growth curves. Mice underwent habituation to head-fixation and learned to associate a visual stimulus with a water reward. Once mice were habituated to head-fixation and associated visual stimuli with water delivery, the mice were initiated on the task.

#### Trial structure

Mice learned and performed a detection task where they reported stimulus detection by licking during a response window (0-1.5 seconds) following stimulus onset. A trial was initiated following a 1-5 second period with no detected licks. Following trial initiation, there was a variable wait time between 1-10 seconds before stimulus presentation. If mice licked before stimulus presentation, the trial was terminated and labeled a “false alarm”. If the trial reached stimulus presentation (i.e., there was no false alarm), the stimulus was presented and in the case of visual stimuli remained on the monitor until a lick was detected or the response window of 1-2 seconds was over. If the mice licked during the response window, the trial was labeled as a “hit” and the mice were given a small water reward (5-20 μL); if the mice did not lick during the response window, the trial was labeled as a “lapse” and no reward was delivered (Figure 8b). “Catch” trials were included which followed the same trial structure, but no stimuli were presented at the onset of the response window. If mice licked during this window, the trial was deemed a “catch false alarm” and was not rewarded.

#### Task design and stimuli

The behavior task featured three phases (Figure 8c). In the initial phase, mice were trained to detect static gratings with varying contrasts (0, 2, 4, 8, 16, 32, 64, 80, and 100% contrasts). During phase 1 training, various methods were employed to facilitate task learning. Mice often developed timing strategies where mice would repetitively lick, resulting in many false alarms. At the trainer’s discretion, triggered by excessive false alarms, these licking strategies were variably negatively punished by enforcing a “false alarm timeout” during which access to the spout was briefly restricted by servo-motor positioning for 3-10 seconds and no trials would be initiated, or positively punished where false alarms would coincide with a mild noxious stimuli including an air puff to the whiskers or a mild electrical shock.

In phase 1, more high-contrast trials were included to promote task learning; however, “test” days were included where all contrasts were included to evaluate detection thresholds. In Phase 2, a new visual stimulus, deemed “moving circle”, was introduced, and moving circle trials were interdispersed with static grating trials. Moving circle stimuli featured circles with varying contrasts (0, 2, 4, 8, 16, 32, 64, 80, and 100% contrasts) and sizes that appeared in random positions on the monitor and had a consistent, slow movement in a random direction. The intention was to replicate the spatially confined, perceptually subtle visual percepts (often referred to as “phosphenes”) reported in human electrical stimulation of V1 studies (cite many options). Mice that demonstrated successful learning of the task in Phase 1 and generalization to the moving circle stimulus in Phase 2 were implanted with stimulating electrodes and proceeded to Phase 3 (see behavior analysis under the subsection “Quantification and Statistical Analyses” for details regarding quantification of performance across phases). In phase 3, static grating trials with varying contrasts, moving circle trials with varying contrasts, and electrical stimuli trials with varying amplitudes were delivered. Generally, electrical stimuli were constrained to single pulses with varying amplitudes, but some sessions specifically tested high-frequency trains (100-300 Hz, with 10-50 pulses) and were analyzed independently.

#### Software and hardware implementation

The behavioral detection task was implemented using custom Python scripts based on the Pyglet library to control stimulus timing, visual presentations, and behavioral events with precise temporal accuracy. Task hardware was interfaced via an Arduino microcontroller (Arduino, MEGA), which managed peripheral devices including an infrared beam sensor (Z4352-ND, DigiKey) for lick detection, a servo motor (2000 Series 5-Turn, Dual Mode Servo 2000-0025-0502, Servocity) for spout positioning, and a solenoid (LHDA1233215H, Lee Co) for water delivery. Visual stimuli were displayed on two adjacent vertical monitors (ASUS VG277Q1A) positioned 30 cm in front of the mice

Detailed documentation of software and hardware, Python task scripts, and parts lists for hardware assembly is available at our public repository (https://github.com/denmanlab/mouse_behavior).

#### Electrophysiology recordings during detection assay

During select behavior sessions, we recorded from contacts not used for stimulation on the chronically implanted 16-contact linear stimulating electrode (Neuronexus, A1x16-5mm-50-703-CM16). In these sessions, two electrode contacts were designated for electrical stimulation while the remaining contacts were routed for electrophysiological recording. The recorded contacts were connected to a RHD 16-Channel Recording Headstages (Intan C3334) with an 18-pin electrode adapter board (Intan C3418) and 18-pin wire adapter (Intan B7600). Neural signals were acquired at 30 kHz using an Open Ephys acquisition system[106]. Post-hoc analysis filtered data into an LFP band (first-order lowpass filtered 300 Hz) and an AP band (first-order highpass filtered 300 Hz). Relevant task timestamps including stimulation onset, reward delivery times, and lick times were recorded on analog and digital lines during the recording.

### Quantification and statistical analysis

#### Dataset

9 mice (5 male, 4 female) were used for acute multi-Neuropixels recordings with electrical stimulation. Single-unit spiking data with associated stimulus information, but without raw binary files, is available in NWB format on DANDI (ID = 000774). Raw binary data can be provided upon request. For training and development of the detection assay, 9 mice were used; four of these mice successfully learned the behavioral task and implanted with a stimulating electrode and ultimately included in the final analyses. The analysis code, including all necessary scripts and intermediates to reproduce the figures, is available at https://github.com/denmanlab/estim_populations.

#### Alignment to CCF coordinates

We used a semi-rigid probe geometry in which probe positions remained consistent relative to one another. Each recording probe was inserted at a 45° angle to the brain surface, and the stimulating electrode was inserted perpendicular to the surface. Recording probes were positioned 100–200 µm horizontally from the stimulating electrode and 100–200 µm vertically from the nearest probe at the crossing point. Probe geometry and relative positioning were recreated using the Neuropixels trajectory planning software [108] to establish trajectories and stereotactic coordinates for surgery. *A priori* CCF coordinates for the Neuropixels probes and stimulating electrode were interpolated from entry and tip coordinates generated in Pinpoint. Before insertion, probe tips were coated with the fluorescent marker CM-DiI for post-hoc histological alignment. Final probe depths were confirmed using electrophysiological features, including LFP power and spike rates.

#### 3D plotting in anatomical brain space

3D plots in CCF mouse anatomical brain space were generated using custom Python code leveraging Brainrender [109](https://github.com/brainglobe/brainrender) and Vedo (https://github.com/marcomusy/vedo). 3D renders with individual single units (Figure 4d, Figure 7e), locations were assigned to the CCf coordinates of the channel with the maximal waveform, with small random mediolateral jitter added for visualization; sphere size represents the amplitude of the single unit waveform.

#### Evoked potential analysis

##### Defining the evoked potential

Evoked potential analyses were performed on raw extracellular voltages from the AP-band. Across all channels in the brain and stimulation conditions, we observed an initial saturating phase – likely due to brief amplifier saturation – that increased with amplitude (Figure 1l, stats) and closely matched the time duration and sign of the biphasic pulse (100 μs for each phase, 200 μs total), suggesting that despite brief saturation, the amplifiers would recover near instantaneously (Figure 2b, example waveform). This saturated artifact is consistent across conditions and varies minimally with distance from the stimulation source, except in rare cases, indicating it does not reflect physiological neural activity. Following this saturation period, we observed a non-saturating waveform component that rapidly returned to baseline due to hardware-imposed highpass filtering in the AP band (Figure 2b). This non-saturating phase scaled with stimulation amplitude and distance from stimulation, supporting its physiological origin rather than being a component of the artifact. This non-saturating phase (Figure 2d, shaded region) was deemed the evoked potential.

##### Measuring evoked potential in AP and LFP band

The evoked potential was recorded in both the AP (acquired at 30,000 samples per second) and LFP band (acquired at 2,500 samples per second). We measured the spatial extent in the AP band, given the much higher sampling precision, allowing more precise measurements of the timepoints immediately preceding stimulation onset. We do, however, note that the hardware highpass filtering (300 Hz) in the AP band would filter slow, persisting induced voltages and thus limit the full temporal profile of the evoked potential. Thus, the analysis of the evoked potential is limited to the spatial extent within the first several milliseconds. This provides additional scientific purpose because measuring the evoked potential within 3 ms of stimulation onset excludes contributions to the electric field that may result from induced synaptic propagation effects at longer timescales.

##### Algorithmically determining the spatial extent of evoked potential

Spatial extent of evoked potentials was algorithmically determined from raw AP band data. Each trial of a given electrical stimulation parameter was aligned to stimulation onset via threshold detection of the saturating portion of the electrical artifact to account for microsecond-level imprecision in synchronized stimulation timestamps. After aligning, a response window from 0.3 ms to 3 ms was selected for each trial to avoid the saturating artifact and formatted into a 3D array (trials, samples, channels), including the first 20 channels out of the brain and in saline. Common signal for a given sample was identified as the median value from the included saline channels and subsequently was subtracted across all channels. The mean across the trials axis was taken, and for further spatial characterization and alignment, samples at 1 ms post-stimulation were analysed (1D array of the trial mean sample at each channel at the 1 ms time bin). For each probe across all recordings, we aligned the 1 ms time bin data to the channel of the maximum voltage, padding the edges when necessary to maintain a homogeneous array shape. This alignment for each probe was then applied to the mean 0.3 - 3 ms window array to create a peak-aligned average evoked potential (samples x channels) for each probe. The evoked response from each probe was averaged to create a mean evoked potential across all probes.

##### Aligning and averaging the evoked potential

To allow spatial averaging across probes, we normalized probe distances by assigning zero microns to the channel showing the peak evoked response. Channels superficial (towards pia) relative to this peak were assigned negative distances, while channels deeper were assigned positive distances, thus enabling consistent spatial comparison across probes. Normalized distances allowed averaging and aligning the evoked potential across probes. However, we transitioned to absolute Euclidean distances derived from CCF coordinates to provide precise anatomical and distance referencing. The trends for distance from stimulation using peak-aligned relative distance (Figure S2e) and Euclidean CCF distance (Figure S2f) were similar. The mean difference in the channel the maximum evoked potential was recorded on and the channel with the minimum Euclidean distance from the CCF coordinates was 8.14 +/- 25.37, corresponding to approximately 80 μm in distance (Figure S2d, i). The difference in channels between peak response and minimum CCF distance did not vary significantly with amplitude (One-way ANOVA: F = 0.883, p = 0.519, Figure S2d, ii), but did vary with recording (One-way ANOVA: F = 23.1621, p < 0.001, Figure S2d, iii). For this normalized distance, every 10 channels towards the surface represented -100 μm and every 10 channels deep to the zero point represented 100 μm. Specific calculations of the spatial extent and magnitude of the evoked potential were performed on voltages at 1 ms. The spatial extent for each direction on a given probe was determined by its return to 20 μV. Area under the curve (AUC) were determined from the boundaries on the evoked potential at 1 ms (Figure S2h).

#### Anisotropic volumetric analyses

##### Volumetric calculations

To calculate volumes, we leveraged the orthogonal probe geometry that sampled the space surrounding the stimulation source. The six total one-dimensional spatial measurements that could be generated from the three Neuropixels (one measurement of the spatial decay superficial to the stimulation source and one deep from the stimulation source on each probe) were used to calculate the volumetric space. We successfully calculated all six spatial extents possible for four of the nine recordings, for all polarities and amplitudes except 5 μA. The CCF coordinates of the six one-dimensional spatial extents were used to fit a convex hull, and we calculated the volume of the convex hull, a conservative estimate of volume. The stimulation site served as the fixed anatomical reference (the “electrode center”). All distances are reported in µm.

##### Isotropic spherical fits

We modeled a symmetric (“isotropic”) evoked-potential profile as a sphere where the center could shift (”free-center” sphere) and where the center was fixed (”fixed-center” sphere) at the source of stimulation. For the free-center sphere, we fit (**c**, r) by nonlinear least squares (SciPy least_squares, with bound r > 0), minimizing the residuals

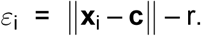

The center shift was:

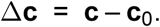

For the fixed center sphere, the radius r_L2_ was defined as the mean Euclidean distance from each point **x**_i_ to the stimulation center **c**_0_:

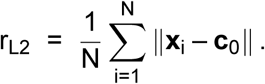

##### Convex-hull comparison to an isotropic sphere hull

Convex hulls summarize the measured 3-D extent but can under-estimate spread along unsampled directions, which the fit spheres do not. To compare the evoked potential hull and the corresponding fit sphere fairly, we:

1. Built the measured (evoked potential) hull from the observed points (SciPy ConvexHull).
2. Constructed a symmetric sphere hull by projecting each measured direction **u**_i_ to the fixed-center sphere with radius r_L2_: **c**_0_ + r_L2_**u**_i_. This preserves the sampling geometry but is isotropic.
3. Quantified continuous deviations by tessellating each triangular face of the measured hull into a fine triangular mesh and computing the signed perpendicular distance of each mesh point to the sphere hull (point-to-triangle distance; sign negative if the point lies inside all sphere-hull half-spaces, positive if outside).

This yields a face-wise field of deviations that we summarize/visualize (coolwarm colormap; units µm). Positive values indicate outward bulging relative to the symmetric sphere; negative values indicate truncation (Figure 3e). Typical tessellation subdivisions per face were n_sub_ = 12–30 (trade-off between smoothness and runtime).

To visualize average condition-level shapes we averaged the CCF coordinates for each spatial extent with a similar probe trajectory and calculated a convex hull from these per-condition averaged coordinates. The condition-average hull is the convex hull of these averaged points placed at the mean stimulation center for that condition. The same sphere-hull and tessellated deviation procedures were then applied to the averaged hull (Figure 3e).

##### Anisotropy metrics

Anisotropy strength (Figure 3f–G).

To quantify directional deviations from a sphere, we modeled the distance of each measured point from the fixed stimulation center as a linear function of its direction. Let **u**_i_ be the unit vector from the fixed center to point i and d_i_ its distance. We fit

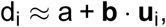

where a is the average radius and **b** captures systematic elongation along a preferred direction. We defined the (dimensionless) anisotropy strength as

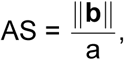

so that AS = 0 indicates a perfect sphere and larger values indicate stronger directional bias.

Center-shift direction metric (Figure 3h).

For each recording, we compared the stimulation center to the center of mass as defined by the mean location of the spatial extents that made up the 3D evoked potential volume. We calculated the direction and magnitude vectors of this shift. Further, we plotted the location of the corpus callosum that was closest to the stimulation site for grounding.

#### Direct response of single units

##### Calculating direct response probability and responders

Direct response probability (DRP) was defined as the proportion of trials in which at least one spike occurred within a 2-ms window following stimulus onset:

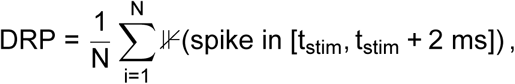

where N is the number of trials and *JJ*(*·*) is the indicator function (equal to 1 if the condition is met, 0 otherwise). To account for baseline firing, a normalized direct response probability (nDRP) was computed by subtracting the probability of spiking in a 2 ms baseline window:

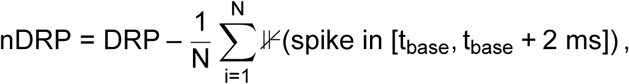

where t_base_ denotes the start of the baseline window. A unit was considered a “responder” to a given stimulus if the normalized DRP exceeded 0.15.

##### Single unit stability metric

Single unit stability (i.e., the ability to record a single unit across the duration of the recording) was essential for the interpretation and accuracy of our results, specifically when comparing responses from different stimulus conditions. To ensure that only stable single units were included in downstream analyses, we quantified firing rate consistency across the duration of each recording session and filtered out unstable units. For each unit, spike times were binned into 60-second non-overlapping intervals across the entire session. Firing rates were calculated as spike count per bin divided by bin duration (Hz). To quantify stability, we fit a linear regression model to each unit’s binned firing rate as a function of time using ordinary least squares (scikit-learn LinearRegression). The resulting slope, intercept, mean squared error (MSE), and coefficient of determination (R²) were recorded. Additionally, we computed the ratio of the y-intercept to the unit’s average firing rate (y-intercept/FR) to assess drift: values near 1 indicate stationarity, while values below or above 1 suggest increasing or decreasing firing rate trends, respectively. Units were excluded if their firing rate trajectory was identified as an outlier using the interquartile range (IQR) method. Specifically, units with either y-intercept/FR falling beyond 1.5 times the IQR from the first or third quartile were labeled as unstable and removed from further analysis. Outlier detection was performed separately for each recording session.

##### Hierarchical clustering of direct responders

We performed hierarchical agglomerative clustering using Ward’s method on normalized feature vectors. The dendrogram was cut into k clusters using the “maxclust” criterion. The clustering quality was evaluated using three complementary metrics: (i) the cophenetic correlation coefficient (CCC), quantifying how well the dendrogram preserved pairwise distances;

**(a)** (ii) the silhouette score, assessing within-cluster cohesion and between-cluster separation; and
**(b)** (iii) the within-cluster sum of squares (WCSS) across different cluster numbers, visualized in an elbow plot.

##### Waveform (FS/RS) classification

Average waveforms were computed for each single unit. Fast-spiking (FS) waveforms, characterized by short, rapidly decaying action potentials, are considered putative parvalbumin-positive (PV+) inhibitory interneurons. Regular spiking (RS) waveforms are characterized by larger, slower waveforms ([52]). Classification was based on spike duration, peak-to-trough ratio, and repolarization slope. Units were initially assigned FS or RS categories by k-means clustering on normalized waveform features. Spikes with positive polarity (peak-to-trough ratio ≥ 1.0) were reclassified as “up” units, and narrow upward spikes (<0.4 ms) were further labeled as putative axonal signals (“axon”), which were not included in analyses unless otherwise stated.

#### Circuit response

##### Determining circuit responders

Circuit responders were determined by applying Zetapy[110] on spike times for a 300 ms window following stimulation onset for each stimulation parameter.

##### Modulation index calculation

We quantified stimulus-evoked changes in firing rate using a Modulation Index (MI). For each unit and trial, the response firing rate was computed from binned spike counts in a 500-1000 ms post-stimulus window using 5 ms bins. The baseline firing rate was defined as the median firing rate over an equivalent pre-stimulus baseline window. The MI was then calculated as:

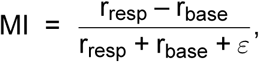

where r_resp_ is the response firing rate, r_base_ is the baseline firing rate, and *ε* is a small constant added to prevent division by zero. This normalization bounds MI between –1 and +1, with positive values indicating higher firing rates during the response window compared to baseline.

##### Dimensionality calculation

We quantified the dimensionality of population activity using the participation ratio (PR) of the eigenvalue spectrum of binned neural firing rates for all cortical neurons, which provides a continuous measure of effective dimensionality. For each time bin (bin size of 2 ms, window of 500 ms), we computed the trial-by-trial covariance matrix across neurons, extracted its eigenvalues, and calculated the PR as:

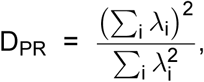

where *λ*_i_ denotes the i^th^ eigenvalue. This metric reflects how many eigenmodes significantly contribute to the variance, with larger values indicating higher-dimensional activity. As a complementary measure, we also computed the minimum number of principal components required to explain 95% of the variance (effective dimensionality). For each time bin, we performed principal component analysis (PCA) on the neuron-by-trial activity matrix, calculated the cumulative explained variance, and defined the effective dimensionality as the smallest number of components exceeding the 95% threshold.

#### Behavior analysis

##### Quantifying task learning

Learning of the task in Phases 1 and 2 was assessed through the calculation of the detection index (d’) for high contrast stimuli. To facilitate behavioral shaping during these initial phases, high contrast stimuli were prioritized, promoting stimulus-response associations. Although many sessions included the full contrast range, there were insufficient catch trials to reliably track performance within individual sessions. Therefore, during Phases 1 and 2, d’ was calculated by leveraging reaction times of rewarded trials for high contrast stimuli, given observations that as mice became proficient, their reaction times for high contrast stimuli narrowed. False positives (F) were operationally defined as trials with reaction times either less than 0.15 s (faster than physiologically possible) or greater than 0.6 s, whereas true positives (H) were defined as rewarded trials with reaction times between 150-600 ms. Thus, d’ was computed as:

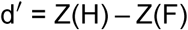

Mice were designated “learned” for phase 1 and 2 stimuli when they experienced a stable d’ measurement over 0 across multiple sessions on high contrast stimuli.

##### Quantifying task performance of electrical and visual stimuli in learned mice

In Phase 3, we evaluated mice that successfully learned to report detection of both visual stimulus classes. In this phase, we included trials with all contrasts of both static gratings and moving circles, and electrical stimulation pulses with varying amplitudes (see “Task design and stimuli”). Notably, sufficient catch trials were introduced, allowing for an accurate assessment of false positive rates during this phase. Here, false positives (F) were defined explicitly as the hit rate on 0% contrast (catch) trials, while true positives (H) were defined as the proportion of rewarded trials at each given contrast or amplitude level.

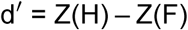

Sessions from Phases 1 and 2 were analyzed to demonstrate task learning and generalization, while all training sessions conducted after successful behavioral shaping (Phase 3) were included in the analysis for determining detection thresholds across stimulus classes. Sessions or intervals characterized by poor task engagement, defined quantitatively by excessive licking without stimulus presentation (false-alarm rate) or failure to respond to high-contrast stimuli (lapse rate), were excluded. Specifically, periods where either the false-alarm rate or lapse rate exceeded 0.7, calculated using a rolling mean over 10 consecutive trials, were omitted from further analyses.

### Data and Code Availability

#### Neurodata Without Borders data format with public availability

Processed data and metadata for each recording were packaged into a Neurodata Without Borders (NWB) format [111] and made available on DANDI Archive (ID: 000774). Metadata included essential experiment and subject metadata including experimenter, electrode information, subject species, sex, and age. Single unit spike times and key features, including but not limited to waveform templates, amplitude, channel, cluster annotations, CCF coordinates, and anatomical regions, were stored in a table structure called “units” in the NWB data structure. Stimulus information including onset times, parameters (including amplitude, polarity, pulse-width, and frequency), and the CCF coordinates for the stimulation contacts were stored in a table structure called “trials” within the NWB data structure where each row represents a stimulation trial. Raw data (LFP band, AP band, and NiDAQ binary files) are not included in the NWB but are available upon request. The NWB for each recording was processed and structured identically, and is in a DANDI-approved format.

#### Analysis code

Analysis code and Python Jupyter notebooks used for analysis and generation of figures and statistics are made available at https://github.com/denmanlab/estim_populations. The purpose of this allows demonstration of all analysis but also independent exploration of data. Python versions and environment details are included. In this repository is also the statistics table.

#### Other code

The code and hardware information used for the behavioral detection task is available at https://github.com/denmanlab/mouse_behavior. The code for Python and Matlab control of the AM-4100 stimulation hardware is available at https://github.com/denmanlab/am4100_code

## Acknowledgements

We thank Juan Santiago-Moreno and Nicholas Garcia for their thoughtful discussions and input. Funding was provided NIH BRAIN Initiative (NINDS) grant R01NS120850 (DJD) and R00EY028612 (DJD).

## Supplemental Figures

**Figure S1.**
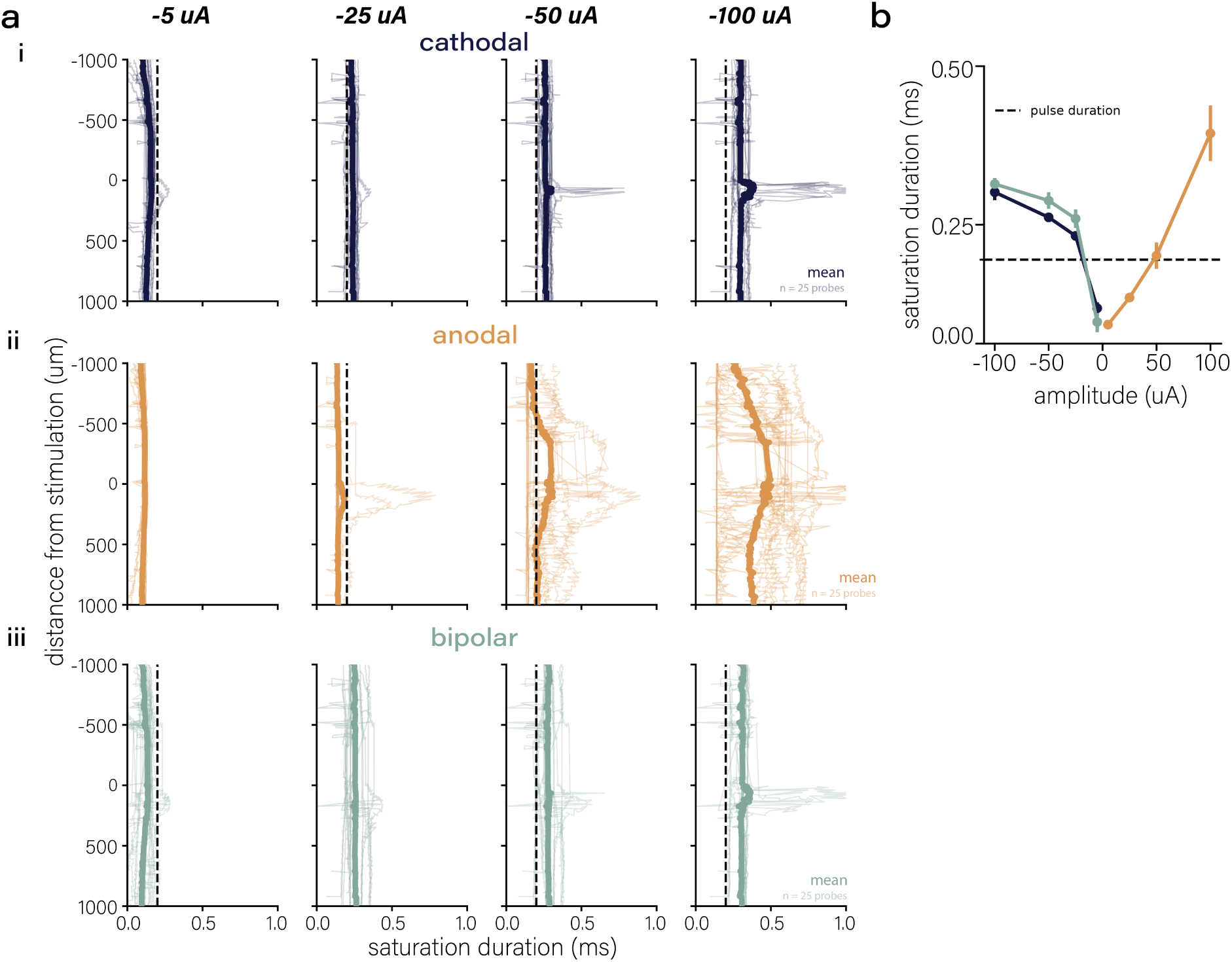
Duration of amplifier saturation following pulses of electrical stimulation across amplitudes and polarities. **(a)** Saturation duration for each cathodal (i), anodal (ii) and bipolar (iii) polarities across distance from stimulation (individual lines represent probes (n=25) **(b)** Average saturation duration across polarities (n = 9 recordings, mean and SEM error clouds).

**Figure S2.**
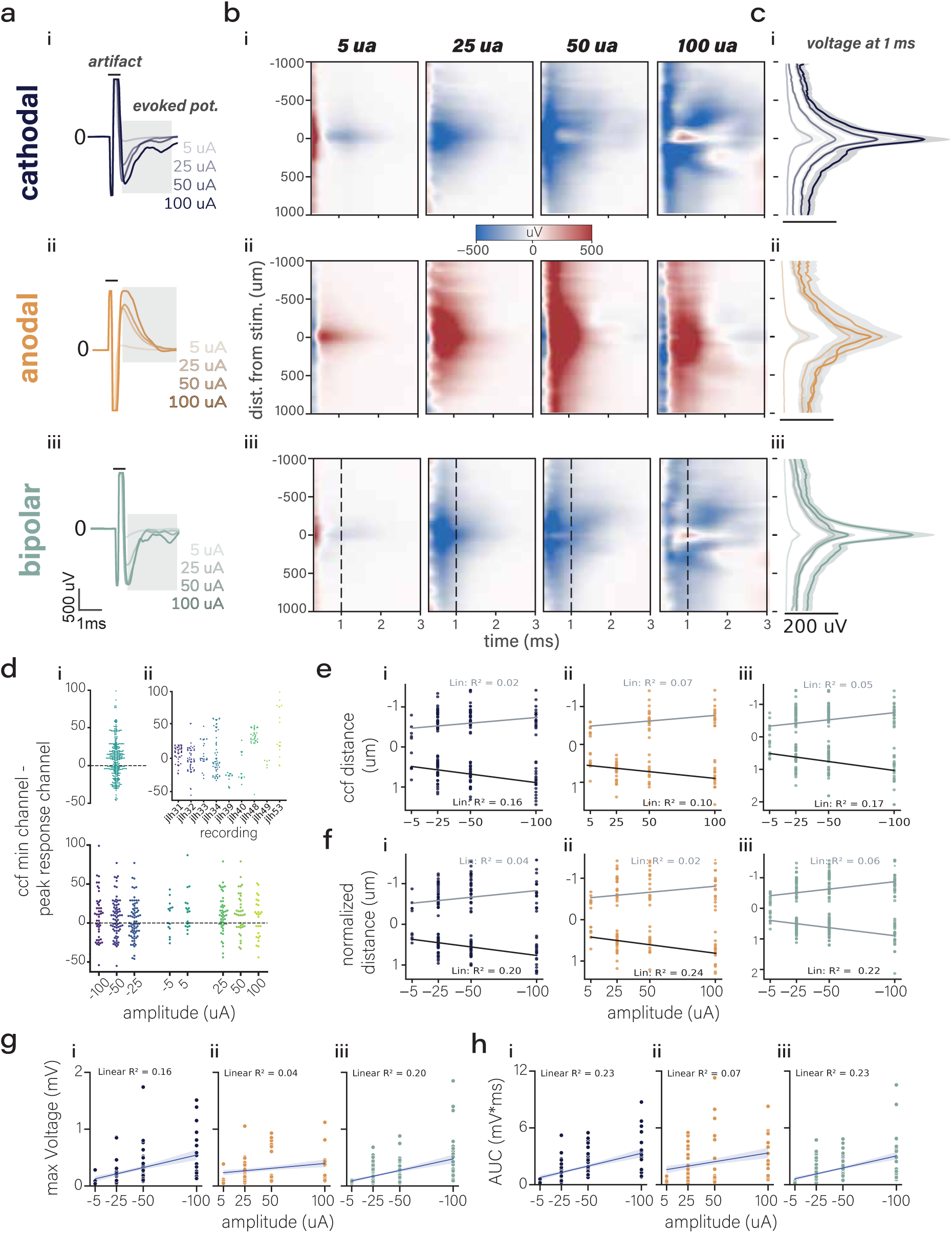
Characterization of evoked potential by polarities. **(a)** average extracellular voltage to single pulses of electrical stimulation of varying amplitudes for cathodal (i), anodal (ii), and bipolar (iii) polarities (n=9 recordings) **(b)** averaged heatmaps aligned to peak response for cathodal (i), anodal (ii), and bipolar (iii) polarities **(c)** average (mean and SEM overlay) extracellular voltage aligned to peak response for cathodal (i), anodal (ii), and bipolar (iii) polarities **(d)** comparison of peak response channel to the channel with the minimum distance from stimulation calculated from CCF coordinates (i) subdivided by recording (ii) and stimulation parameter (iii) **(e)** linear regressions for distance from stimulation at 1 ms using absolute CCF distance as a function of amplitude for cathodal (i), anodal (ii), and bipolar (iii) polarities **(f)** linear regressions for distance from stimulation at 1 ms using peak aligned distance as a function of amplitude for cathodal (i), anodal (ii), and bipolar (iii) polarities **(g)** linear regressions for the maximum voltages at 1 ms as a function of amplitude for cathodal (i), anodal (ii), and bipolar (iii) polarities **(h)** linear regressions for area under the curve at 1 ms as a function of amplitude for cathodal (i), anodal (ii), and bipolar (iii) polarities.

**Figure S3.**
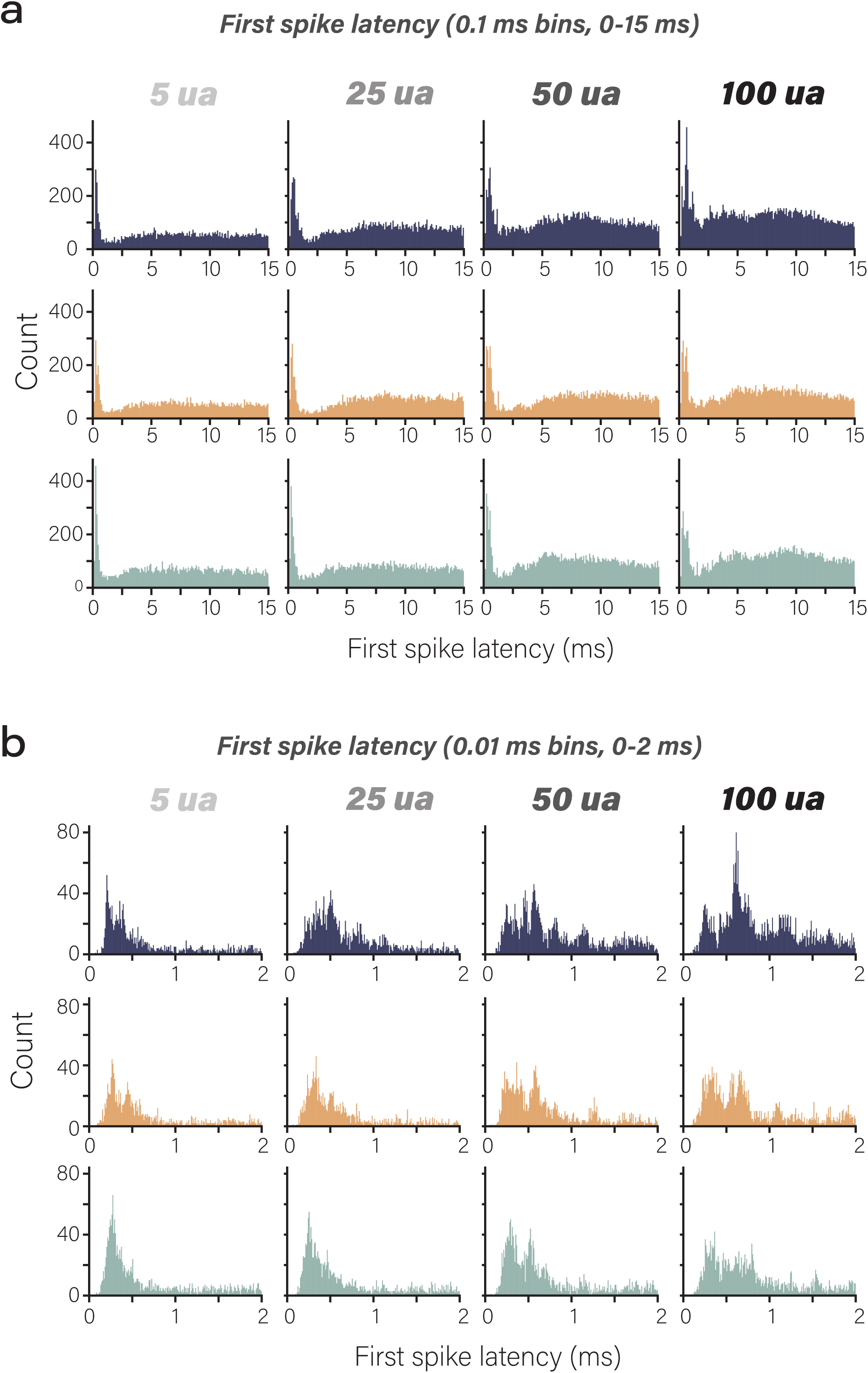
Spike latencies from stimulation by amplitude and polarity. **(a)** Spike latencies of a 15 ms window (0.1 ms bin width) following stimulation onset for varying amplitudes (columns) and polarity (rows) **(b)** Spike latencies of a 2 ms window (0.01 ms bin width) following stimulation onset for varying amplitudes (columns) and polarity (rows).

**Figure S4.**
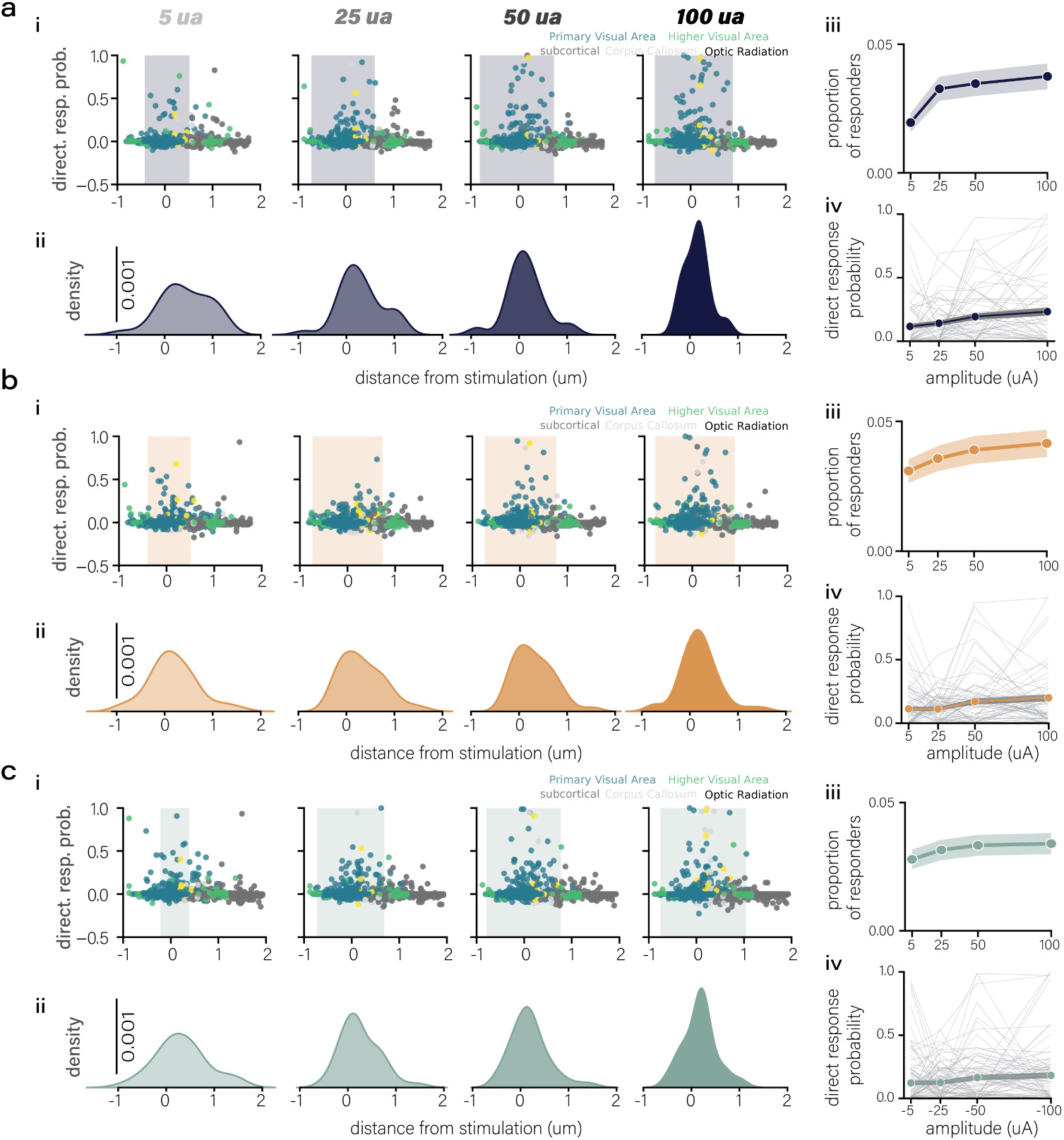
Direct response features by polarities and amplitudes. **(a)** cathodal: direct response probability (i) and responder density (ii) as a function of distance from stimulation. iii, proportion of responders by amplitude. iv, direct response probability of responders by amplitude **(b)** anodal: direct response probability (i) and responder density (ii) as a function of distance from stimulation. iii, proportion of responders by amplitude. iv, direct response probability of responders by amplitude **(c)** bipolar: direct response probability (i) and responder density (ii) as a function of distance from stimulation. iii, proportion of responders by amplitude. iv, direct response probability of responders by amplitude.

**Figure S5.**
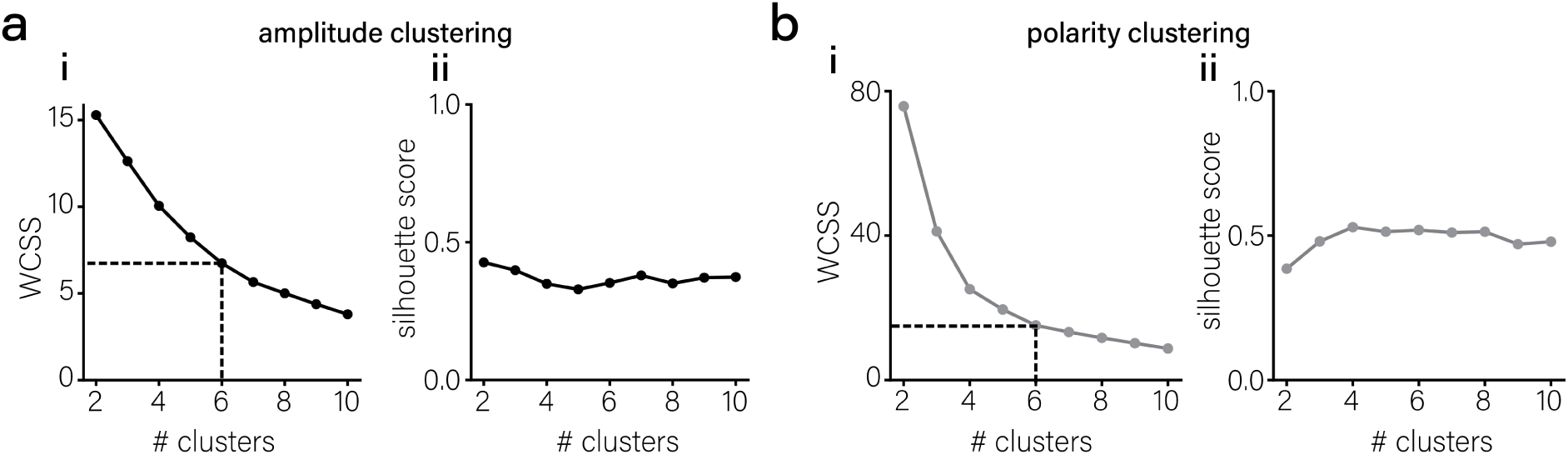
Spatial distribution of direct response for FS and RS single units by polarity. **(a)** clustering fit statistics for amplitude clustering (see Figure 5b). i, the sum of the variance between the observations in each cluster (WCSS). ii, silhouette score for each cluster **(b)** clustering fit statistics for polarity clustering (see Figure 5e). i, the sum of the variance between the observations in each cluster (WCSS). ii, silhouette score for each cluster.

**Figure S6.**
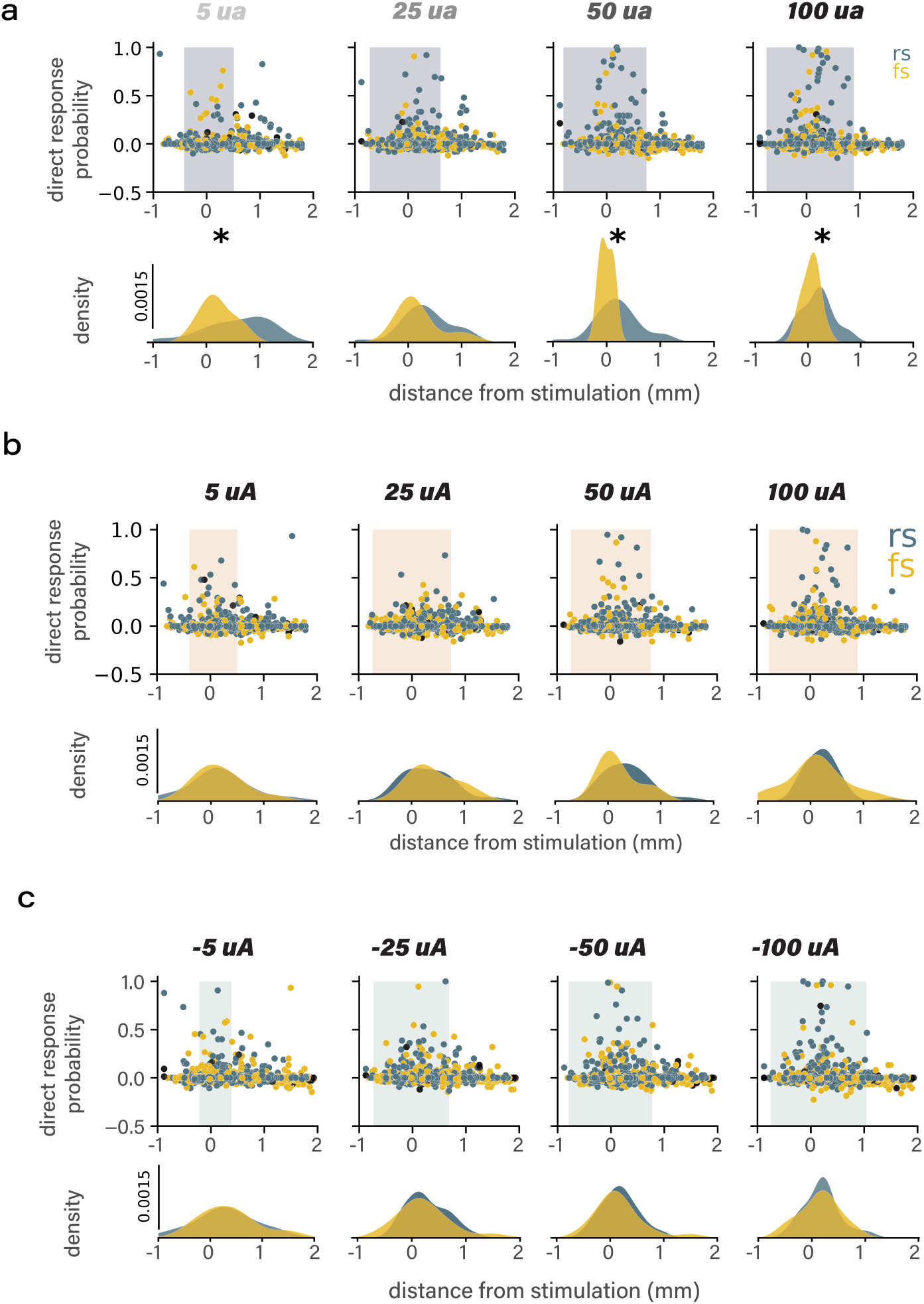
Spatial distribution of direct response for FS and RS single units by polarity. **(a)** cathodal direct response probability (top) and responder density (bottom) for FS and RS single units **(b)** anodal direct response probability (top) and responder density (bottom) for FS and RS single units **(c)** bipolar direct response probability (top) and responder density (bottom) for FS and RS single units.

**Figure S7.**
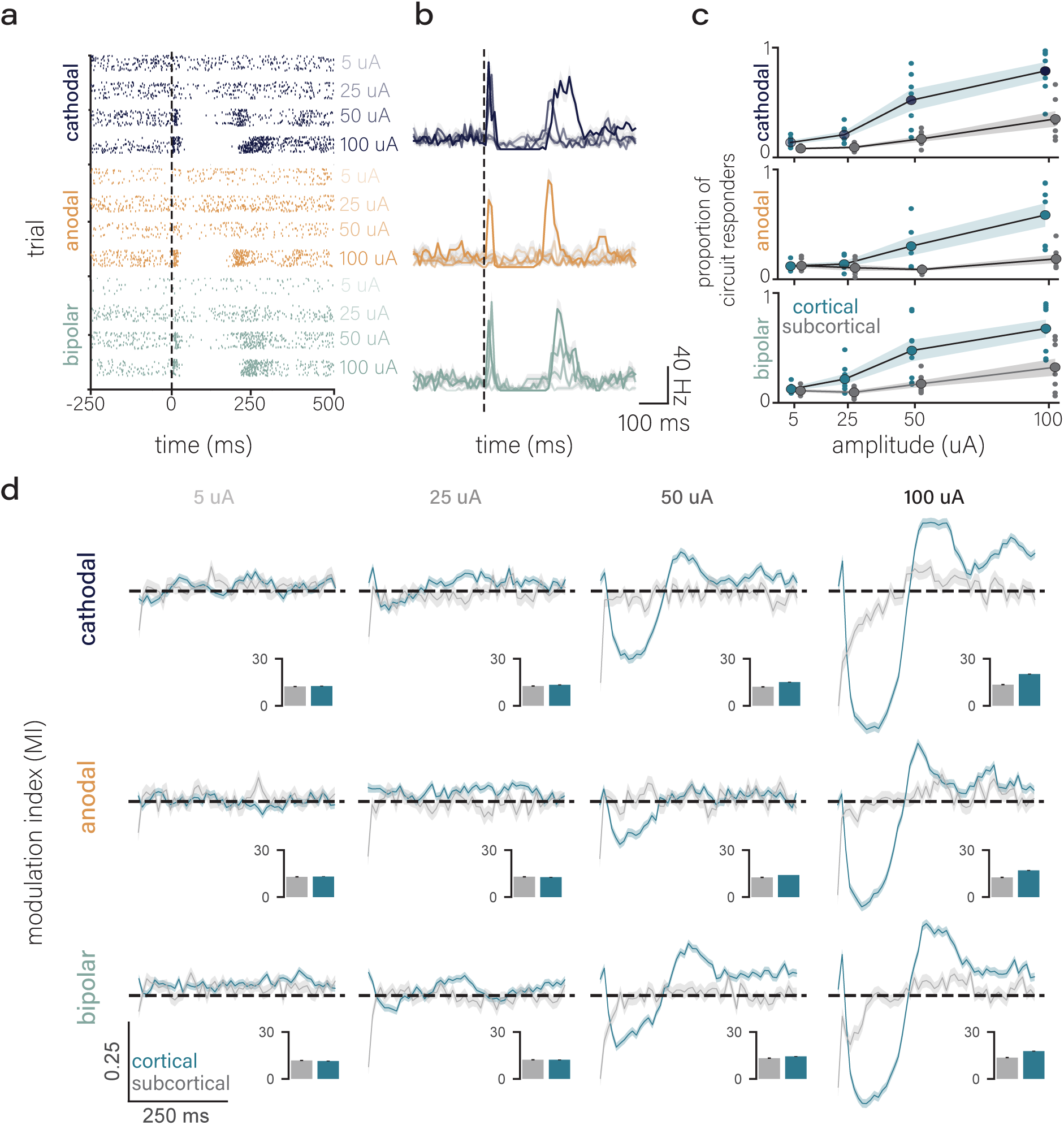
Circuit response by amplitude and polarity. **(a)** rasters for example single unit for each amplitude and polarity **(b)** averaged (mean and SEM) firing rate for an example neuron by amplitude and polarity **(c)** proportion of responders as a function of distance from stimulation for each polarity (mean and SEM overlay) **(d)** modulation index (MI) for a 500 ms window aligned to stimulation onset for each amplitude and polarity (mean and SEM cloud, n = 9 recordings).

**Figure S8.**
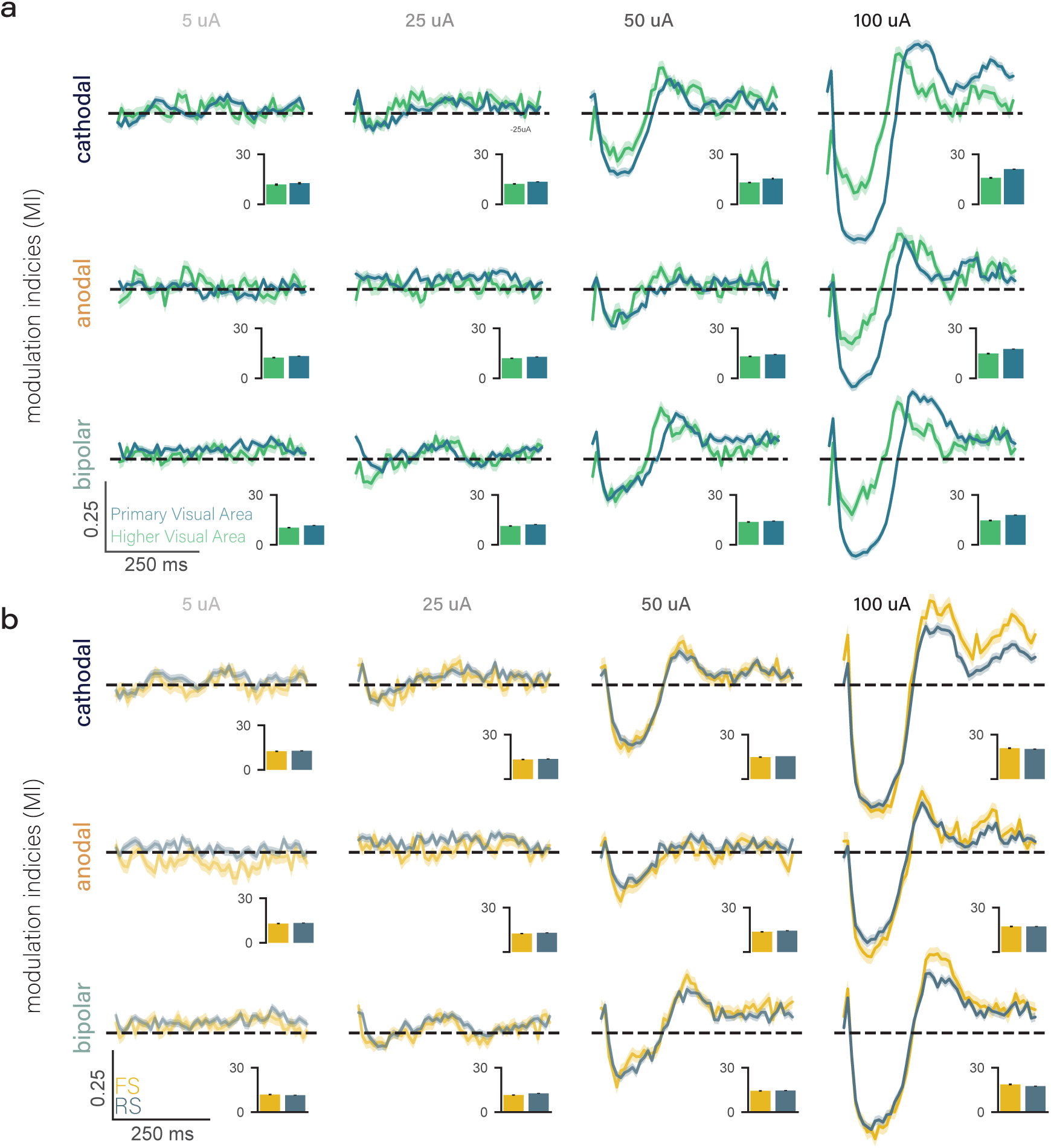
Modulation indices (MIs) by amplitude and polarity for brain regions and single unit waveform class. **(a)** MIs for primary visual area (blue) and higher visual areas (green) for cathodal (top row), anodal (middle row) and bipolar (bottom row) and amplitudes (columns). (mean lines and SEM clouds, bin size = 5 ms, n = 9 recordings). Inset: cumulative modulation **(b)** MIs for FS (yellow) and RS (blue) single units for cathodal (top row), anodal (middle row) and bipolar (bottom row) and amplitudes (columns). (mean lines and SEM clouds, bin size = 5 ms, n = 9 recordings). Inset: cumulative modulation **(c)** Normalized participation ratio (left y-axis, blue) and normalized firing right (right y-axis, black) across amplitudes for cathodal polarity (mean lines and SEM clouds, bin size = 5 ms, n = 9 recordings).

**Figure S9.**
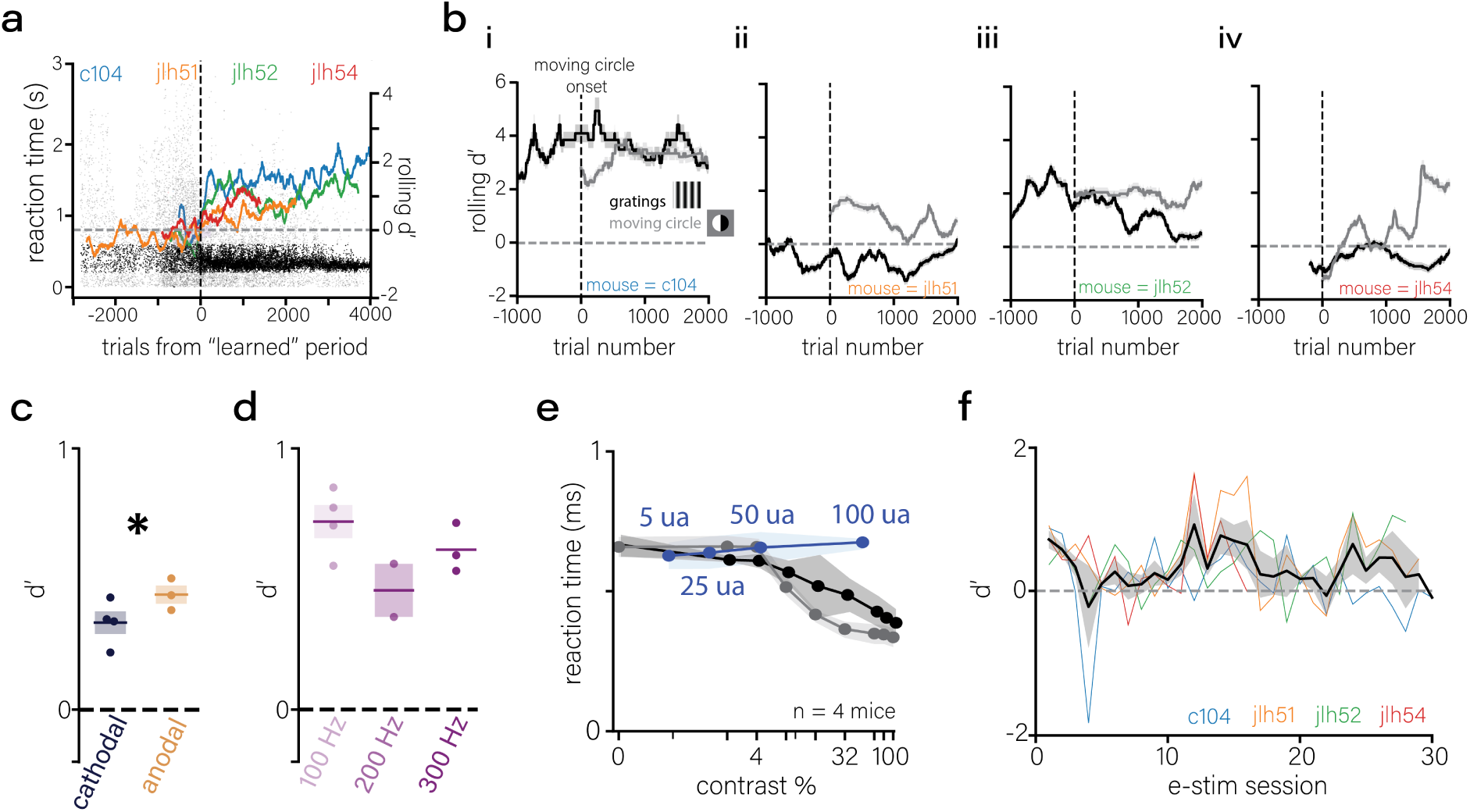
Additional behavioral metrics for detection assay. **(a)** Reaction times (scatter, left axis) and rolling d′ (lines, right axis) for high-contrast (>30%) gratings in mice advancing to Phase 3. Black points: rewarded trials with reaction times 0.2–0.6 s (true positives during learning). Gray points: reaction times <0.2 s or >0.6 s (false positives) included in d′ calculation **(b)** rolling d’ for static gratings and moving circle visual stimuli aligned to the introduction of the moving circle task for each mouse (i-iv) **(c)** d’ for cathodal and anodal pulses (n = 4, mean line and SEM error, paired T-test, p = 0.028) **(d)** d’ for different frequencies (n = 4, mean line and SEM error, One-way ANOVA, p = 0.112) **(e)** reaction times for static gratings (black), moving circle (gray), and electrical stimulation (blue) **(f)** d’ for electrical stimulation trials across sessions in Phase 3 (black line and shaded region represent mean and SEM, individual colors represent mice, n = 4).

## References

1. Salzman, C. D., Britten, K. H., and Newsome, W. T. (1990). Cortical microstimulation influences perceptual judgements of motion direction. en. Nature 346. Publisher: Nature Publishing Group, 174–177. 10.1038/346174a0.

2. Graziano, M. S. A., Taylor, C. S. R., and Moore, T. (2002). Complex Movements Evoked by Microstimulation of Precentral Cortex. Neuron 34, 841–851. 10.1016/S0896-6273(02)00698-0.

3. Afraz, S.-R., Kiani, R., and Esteky, H. (2006). Microstimulation of inferotemporal cortex influences face categorization. en. Nature 442. Publisher: Nature Publishing Group, 692– 695. 10.1038/nature04982.

4. Benabid, A., Pollak, P., Louveau, A., Henry, S., and Rougemont, J. de (1988). Combined (Thalamotomy and Stimulation) Stereotactic Surgery of the VIM Thalamic Nucleus for Bilateral Parkinson Disease. Applied Neurophysiology 50, 344–346. 10.1159/000100803.

5. Benabid, A. L., Pollak, P., Gervason, C., Hoffmann, D., Gao, D. M., Hommel, M., Perret, J. E., and Rougemont, J. de (1991). Long-term suppression of tremor by chronic stimulation of the ventral intermediate thalamic nucleus. eng. Lancet (London, England) 337, 403–406. 10.1016/0140-6736(91)91175-t.

6. Hickey, P. and Stacy, M. (2016). Deep Brain Stimulation: A Paradigm Shifting Approach to Treat Parkinson’s Disease. english. Frontiers in Neuroscience 10. Publisher: Frontiers. 10.3389/fnins.2016.00173.

7. Lyons, K. E. and Pahwa, R. (2008). Deep Brain Stimulation and Tremor. Neurotherapeutics. Movement Disorders 5, 331–338. 10.1016/j.nurt.2008.01.004.

8. Hu, W. and Stead, M. (2014). Deep brain stimulation for dystonia. Translational Neurodegeneration 3, 2. 10.1186/2047-9158-3-2.

9. Mayberg, H. S., Lozano, A. M., Voon, V., McNeely, H. E., Seminowicz, D., Hamani, C., Schwalb, J. M., and Kennedy, S. H. (2005). Deep Brain Stimulation for Treatment-Resistant Depression. Neuron 45, 651–660. 10.1016/j.neuron.2005.02.014.

10. Mar-Barrutia, L., Real, E., Segalás, C., Bertolín, S., Menchón, J. M., and Alonso, P. (2021). Deep brain stimulation for obsessive-compulsive disorder: A systematic review of worldwide experience after 20 years. World Journal of Psychiatry 11, 659–680. 10.5498/wjp.v11.i9.659.

11. Corripio, I., Roldán, A., Sarró, S., McKenna, P. J., Alonso-Solís, A., Rabella, M., Díaz, A., Puigdemont, D., Pérez-Solà, V., Álvarez, E., et al. (2020). Deep brain stimulation in treatment resistant schizophrenia: A pilot randomized cross-over clinical trial. EBioMedicine 51, 102568. 10.1016/j.ebiom.2019.11.029.

12. Boon, P., Raedt, R., Herdt, V. de, Wyckhuys, T., and Vonck, K. (2009). Electrical stimulation for the treatment of epilepsy. Neurotherapeutics 6, 218–227. 10.1016/j.nurt.2008.12.003.

13. Armenta Salas, M., Bashford, L., Kellis, S., Jafari, M., Jo, H., Kramer, D., Shanfield, K., Pejsa, K., Lee, B., Liu, C. Y., et al. (2018). Proprioceptive and cutaneous sensations in humans elicited by intracortical microstimulation. eLife 7. Ed. by R. Romo. Publisher: eLife Sciences Publications, Ltd, e32904. 10.7554/eLife.32904.

14. Quick, K. M., Weiss, J. M., Clemente, F., Gaunt, R. A., and Collinger, J. L. (2020). Intracortical Microstimulation Feedback Improves Grasp Force Accuracy in a Human Using a Brain-Computer Interface. Annual International Conference of the IEEE Engineering in Medicine and Biology Society. IEEE Engineering in Medicine and Biology Society. Annual International Conference 2020, 3355–3358. 10.1109/EMBC44109.2020.9175926.

15. Flesher, S. N., Collinger, J. L., Foldes, S. T., Weiss, J. M., Downey, J. E., Tyler-Kabara, E. C., Bensmaia, S. J., Schwartz, A. B., Boninger, M. L., and Gaunt, R. A. (2016). Intracortical microstimulation of human somatosensory cortex. Science Translational Medicine 8. Publisher: American Association for the Advancement of Science, 361ra141–361ra141. 10.1126/scitranslmed.aaf8083.

16. Vugt, B. van, Dagnino, B., Vartak, D., Safaai, H., Panzeri, S., Dehaene, S., and Roelfsema, P. R. (2018). The threshold for conscious report: Signal loss and response bias in visual and frontal cortex. Science 360. Publisher: American Association for the Advancement of Science, 537–542. 10.1126/science.aar7186.

17. Flesher, S. N., Downey, J. E., Weiss, J. M., Hughes, C. L., Herrera, A. J., Tyler-Kabara, E. C., Boninger, M. L., Collinger, J. L., and Gaunt, R. A. (2021). A brain-computer interface that evokes tactile sensations improves robotic arm control. Science 372. Publisher: American Association for the Advancement of Science, 831–836. 10.1126/science.abd0380.

18. Chen, X., Wang, F., Fernandez, E., and Roelfsema, P. R. (2020). Shape perception via a high-channel-count neuroprosthesis in monkey visual cortex. Science 370. Publisher: American Association for the Advancement of Science, 1191–1196. 10.1126/science.abd7435.

19. Klink, P. C., Dagnino, B., Gariel-Mathis, M.-A., and Roelfsema, P. R. (2017). Distinct Feedforward and Feedback Effects of Microstimulation in Visual Cortex Reveal Neural Mechanisms of Texture Segregation. Neuron 95, 209–220.e3. 10.1016/j.neuron.2017.05.033.

20. Fernández, E., Alfaro, A., Soto-Sánchez, C., Gonzalez-Lopez, P., Lozano, A. M., Peña, S., Grima, M. D., Rodil, A., Gómez, B., Chen, X., et al. (2021). Visual percepts evoked with an intracortical 96-channel microelectrode array inserted in human occipital cortex. en. The Journal of Clinical Investigation 131. Publisher: American Society for Clinical Investigation. 10.1172/JCI151331.

21. Hughes, C. L., Flesher, S. N., and Gaunt, R. A. (2022). EFFECTS OF STIMULUS PULSE RATE ON SOMATOSENSORY ADAPTATION IN THE HUMAN CORTEX. Brain stimulation 15, 987–995. 10.1016/j.brs.2022.05.021.

22. Greenspon, C. M., Valle, G., Shelchkova, N. D., Hobbs, T. G., Verbaarschot, C., Callier, T., Berger-Wolf, E. I., Okorokova, E. V., Hutchison, B. C., Dogruoz, E., et al. (2025). Evoking stable and precise tactile sensations via multi-electrode intracortical microstimulation of the somatosensory cortex. en. Nature Biomedical Engineering 9. Publisher: Nature Publishing Group, 935–951. 10.1038/s41551-024-01299-z.

23. Brindley, G. S. and Lewin, W. S. (1968). The sensations produced by electrical stimulation of the visual cortex. The Journal of Physiology 196, 479–493.

24. Dobelle, W. H. and Mladejovsky, M. G. (1974). Phosphenes produced by electrical stimulation of human occipital cortex, and their application to the development of a prosthesis for the blind. The Journal of Physiology 243, 553–576.1.

25. Dobelle, W. H., Mladejovsky, M. G., Evans, J. R., Roberts, T. S., and Girvin, J. P. (1976). ”Braille” reading by a blind volunteer by visual cortex stimulation. eng. Nature 259, 111–112. 10.1038/259111a0.

26. Dobelle, W. H. (2000). Artificial vision for the blind by connecting a television camera to the visual cortex. eng. ASAIO journal (American Society for Artificial Internal Organs: 1992) 46, 3–9. 10.1097/00002480-200001000-00002.

27. Lewis, P. M. and Rosenfeld, J. V. (2016). Electrical stimulation of the brain and the development of cortical visual prostheses: An historical perspective. eng. Brain Research 1630, 208–224. 10.1016/j.brainres.2015.08.038.

28. Tehovnik, E. J., Slocum, W. M., Smirnakis, S. M., and Tolias, A. S. (2009). Microstimulation of visual cortex to restore vision. eng. Progress in Brain Research 175, 347–375. 10.1016/S0079-6123(09)17524-6.

29. Fernandez, E. and Robles, J. A. (2024). Advances and challenges in the development of visual prostheses. PLOS Biology 22, e3002896. 10.1371/journal.pbio.3002896.

30. Oswalt, D., Bosking, W., Sun, P., Sheth, S. A., Niketeghad, S., Salas, M. A., Patel, U., Greenberg, R., Dorn, J., Pouratian, N., et al. (2021). Multi-electrode stimulation evokes consistent spatial patterns of phosphenes and improves phosphene mapping in blind subjects. en. Brain Stimulation 14, 1356–1372. 10.1016/j.brs.2021.08.024.

31. Urdaneta, M. E., Kunigk, N. G., Delgado, F., Fried, S. I., and Otto, K. J. (2021). Layer-specific parameters of intracortical microstimulation of the somatosensory cortex. en. Journal of Neural Engineering 18. Publisher: IOP Publishing, 055007. 10.1088/1741-2552/abedde.

32. Kunigk, N. G., Urdaneta, M. E., Malone, I. G., Delgado, F., and Otto, K. J. (2022). Reducing Behavioral Detection Thresholds per Electrode via Synchronous, Spatially-Dependent Intracortical Microstimulation. eng. Frontiers in Neuroscience 16, 876142. 10.3389/fnins.2022.876142.

33. Beauchamp, M. S., Oswalt, D., Sun, P., Foster, B. L., Magnotti, J. F., Niketeghad, S., Pouratian, N., Bosking, W. H., and Yoshor, D. (2020). Dynamic Stimulation of Visual Cortex Produces Form Vision in Sighted and Blind Humans. Cell 181, 774–783.e5. 10.1016/j.cell.2020.04.033.

34. Greenspon, C. M., Shelchkova, N. D., Valle, G., Hobbs, T. G., Berger-Wolf, E. I., Hutchison, B. C., Dogruoz, E., Verbarschott, C., Callier, T., Sobinov, A. R., et al. (2023). Tessellation of artificial touch via microstimulation of human somatosensory cortex. bioRxiv, 2023.06.23.54542 10.1101/2023.06.23.545425.

35. Valle, G., Alamri, A. H., Downey, J. E., Lienkämper, R., Jordan, P. M., Sobinov, A. R., Endsley, L. J., Prasad, D., Boninger, M. L., Collinger, J. L., et al. (2025). Tactile edges and motion via patterned microstimulation of the human somatosensory cortex. Science 387. Publisher: American Association for the Advancement of Science, 315–322. 10.1126/science.adq5978.

36. Ranck, J. B. (1975). Which elements are excited in electrical stimulation of mammalian central nervous system: A review. en. Brain Research 98, 417–440. 10.1016/0006-8993(75)90364-9.

37. Stoney, S. D., Thompson, W. D., and Asanuma, H. (1968). Excitation of pyramidal tract cells by intracortical microstimulation: effective extent of stimulating current. Journal of Neurophysiology 31. Publisher: American Physiological Society, 659–669. 10.1152/jn.1968.31.5.659.

38. Histed, M. H., Bonin, V., and Reid, R. C. (2009). Direct activation of sparse, distributed populations of cortical neurons by electrical microstimulation. eng. Neuron 63, 508–522. 10.1016/j.neuron.2009.07.016.

39. Kumaravelu, K., Sombeck, J., Miller, L. E., Bensmaia, S. J., and Grill, W. M. (2022). Stoney vs. Histed: Quantifying the spatial effects of intracortical microstimulation. Brain Stimulation 15, 141–151. 10.1016/j.brs.2021.11.015.

40. Jun, J. J., Steinmetz, N. A., Siegle, J. H., Denman, D. J., Bauza, M., Barbarits, B., Lee, A. K., Anastassiou, C. A., Andrei, A., Aydın, Ç., et al. (2017). Fully integrated silicon probes for high-density recording of neural activity. en. Nature 551. Publisher: Nature Publishing Group, 232–236. 10.1038/nature24636.

41. Wang, Q., Ding, S.-L., Li, Y., Royall, J., Feng, D., Lesnar, P., Graddis, N., Naeemi, M., Facer, B., Ho, A., et al. (2020). The Allen Mouse Brain Common Coordinate Framework: A 3D Reference Atlas. Cell 181, 936–953.e20. 10.1016/j.cell.2020.04.007.

42. McIntyre, C. C. and Grill, W. M. (2001). Finite element analysis of the current-density and electric field generated by metal microelectrodes. eng. Annals of Biomedical Engineering 29, 227–235. 10.1114/1.1352640.

43. — (2000). Selective microstimulation of central nervous system neurons. eng. Annals of Biomedical Engineering 28, 219–233. 10.1114/1.262.

44. Li, C.-l., Bak, A. F., and Parker, L. O. (1968). Specific resistivity of the cerebral cortex and white matter. Experimental Neurology 20, 544–557. 10.1016/0014-4886(68)90108-8.

45. Ranck, J. B. (1963). Specific impedance of rabbit cerebral cortex. eng. Experimental Neurology 7, 144–152. 10.1016/s0014-4886(63)80005-9.

46. Brocker, D. T. and Grill, W. M. (2013). “Chapter 1 - Principles of electrical stimulation of neural tissue”. Handbook of Clinical Neurology. Ed. by A. M. Lozano and M. Hallett. Vol. 116. Brain Stimulation. Elsevier, 3–18. 10.1016/B978-0-444-53497-2.00001-2.

47. Gorman, A. L. (1966). Differential patterns of activation of the pyramidal system elicited by surface anodal and cathodal cortical stimulation. eng. Journal of Neurophysiology 29, 547–564. 10.1152/jn.1966.29.4.547.

48. Grill, W. and Mortimer, J. (1995). Stimulus waveforms for selective neural stimulation. IEEE Engineering in Medicine and Biology Magazine 14, 375–385. 10.1109/51.395310.

49. McIntyre, C. C., Grill, W. M., Sherman, D. L., and Thakor, N. V. (2004). Cellular effects of deep brain stimulation: model-based analysis of activation and inhibition. eng. Journal of Neurophysiology 91, 1457–1469. 10.1152/jn.00989.2003.

50. Butovas, S. and Schwarz, C. (2003). Spatiotemporal Effects of Microstimulation in Rat Neocortex: A Parametric Study Using Multielectrode Recordings. Journal of Neurophysiology 90. Publisher: American Physiological Society, 3024–3039. 10.1152/jn.00245.2003.

51. Hao, Y., Riehle, A., and Brochier, T. G. (2016). Mapping Horizontal Spread of Activity in Monkey Motor Cortex Using Single Pulse Microstimulation. Frontiers in Neural Circuits 10, 104. 10.3389/fncir.2016.00104.

52. Barthó, P., Hirase, H., Monconduit, L., Zugaro, M., Harris, K. D., and Buzsáki, G. (2004). Characterization of Neocortical Principal Cells and Interneurons by Network Interactions and Extracellular Features. Journal of Neurophysiology 92. Publisher: American Physiological Society, 600–608. 10.1152/jn.01170.2003.

53. Logothetis, N. K., Augath, M., Murayama, Y., Rauch, A., Sultan, F., Goense, J., Oeltermann, A., and Merkle, H. (2010). The effects of electrical microstimulation on cortical signal propagation. en. Nature Neuroscience 13. Publisher: Nature Publishing Group, 1283–1291. 10.1038/nn.2631.

54. Yun, R., Mishler, J. H., Perlmutter, S. I., Rao, R. P. N., and Fetz, E. E. (2023). Responses of Cortical Neurons to Intracortical Microstimulation in Awake Primates. en. eNeuro 10. Publisher: Society for Neuroscience Section: Research Article: New Research. 10.1523/ENEURO.0336-22.2023.

55. Hajnal, B., Szabó, J. P., Tóth, E., Keller, C. J., Wittner, L., Mehta, A. D., Erõss, L., Ulbert, I., Fabó, D., and Entz, L. (2024). Intracortical mechanisms of single pulse electrical stimulation (SPES) evoked excitations and inhibitions in humans. en. Scientific Reports 14. Publisher: Nature Publishing Group, 13784. 10.1038/s41598-024-62433-0.

56. O’Rawe, J. F., Zhou, Z., Li, A. J., LaFosse, P. K., Goldbach, H. C., and Histed, M. H. (2023). Excitation creates a distributed pattern of cortical suppression due to varied recurrent input. english. Neuron 111. Publisher: Elsevier, 4086–4101.e5. 10.1016/j.neuron.2023.09.010.

57. Cruikshank, S. J., Lewis, T. J., and Connors, B. W. (2007). Synaptic basis for intense thalamocortical activation of feedforward inhibitory cells in neocortex. eng. Nature Neuroscience 10, 462–468. 10.1038/nn1861.

58. Cruikshank, S. J., Urabe, H., Nurmikko, A. V., and Connors, B. W. (2010). Pathway-Specific Feedforward Circuits between Thalamus and Neocortex Revealed by Selective Optical Stimulation of Axons. english. Neuron 65. Publisher: Elsevier, 230–245. 10.1016/j.neuron.2009.12.025.

59. Stringer, C., Pachitariu, M., Steinmetz, N., Carandini, M., and Harris, K. D. (2019). High-dimensional geometry of population responses in visual cortex. en. Nature 571, 361–365. 10.1038/s41586-019-1346-5.

60. Cone, J. J., Ni, A. M., Ghose, K., and Maunsell, J. H. R. (2018). Electrical Microstimulation of Visual Cerebral Cortex Elevates Psychophysical Detection Thresholds. eNeuro 5, ENEURO.0311–18.2018. 10.1523/ENEURO.0311-18.2018.

61. Beyeler, M., Rokem, A., Boynton, G. M., and Fine, I. (2017). Learning to see again: Biological constraints on cortical plasticity and the implications for sight restoration technologies. Journal of neural engineering 14, 051003. 10.1088/1741-2552/aa795e.

62. Fricke, H. (1953). The Electric Permittivity of a Dilute Suspension of Membrane-Covered Ellipsoids. Journal of Applied Physics 24, 644–646. 10.1063/1.1721343.

63. Tehovnik, E. J. (1996). Electrical stimulation of neural tissue to evoke behavioral responses. eng. Journal of Neuroscience Methods 65, 1–17. 10.1016/0165-0270(95)00131-x.

64. Cushing, H. (1909). A NOTE UPON THE FARADIC STIMULATION OF THE POST-CENTRAL GYRUS IN CONSCIOUS PATIENTS.1. Brain 32, 44–53. 10.1093/brain/32.1.44.

65. Limousin, P., Pollak, P., Benazzouz, A., Hoffmann, D., Le Bas, J. F., Broussolle, E., Perret, J. E., and Benabid, A. L. (1995). Effect of parkinsonian signs and symptoms of bilateral subthalamic nucleus stimulation. eng. Lancet (London, England) 345, 91–95. 10.1016/s0140-6736(95)90062-4.

66. Rattay, F. (1986). Analysis of Models for External Stimulation of Axons. IEEE Transactions on Biomedical Engineering BME*-*33, 974–977. 10.1109/TBME.1986.325670.

67. Rattay, F. (1999). The basic mechanism for the electrical stimulation of the nervous system. eng. Neuroscience 89, 335–346. 10.1016/s0306-4522(98)00330-3.

68. McIntyre, C. C., Richardson, A. G., and Grill, W. M. (2002). Modeling the Excitability of Mammalian Nerve Fibers: Influence of Afterpotentials on the Recovery Cycle. Journal of Neurophysiology 87. Publisher: American Physiological Society, 995–1006. 10.1152/jn.00353.2001.

69. Margalit, S. N. and Slovin, H. (2018). Spatio-temporal characteristics of population responses evoked by microstimulation in the barrel cortex. en. Scientific Reports 8. Number: 1 Publisher: Nature Publishing Group, 13913. 10.1038/s41598-018-32148-0.

70. Tehovnik, E. J., Tolias, A. S., Sultan, F., Slocum, W. M., and Logothetis, N. K. (2006). Direct and indirect activation of cortical neurons by electrical microstimulation. eng. Journal of Neurophysiology 96, 512–521. 10.1152/jn.00126.2006.

71. Rattay, F. and Wenger, C. (2010). Which elements of the mammalian central nervous system are excited by low current stimulation with microelectrodes? Neuroscience 170, 399–407. 10.1016/j.neuroscience.2010.07.032.

72. Kole, M. H. P., Ilschner, S. U., Kampa, B. M., Williams, S. R., Ruben, P. C., and Stuart, G. J. (2008). Action potential generation requires a high sodium channel density in the axon initial segment. en. Nature Neuroscience 11. Publisher: Nature Publishing Group, 178–186. 10.1038/nn2040.

73. Hu, W., Tian, C., Li, T., Yang, M., Hou, H., and Shu, Y. (2009). Distinct contributions of Na(v)1.6 and Na(v)1.2 in action potential initiation and backpropagation. eng. Nature Neuroscience 12, 996–1002. 10.1038/nn.2359.

74. Garcia, J. D., Wang, C., Alexander, R. P., Banks, E., Fenton, T., DeKeyser, J.-M., Abramova, T. V., George Jr, A. L., Ben-Shalom, R., Hackos, D. H., et al. (2025). Differential roles of NaV1.2 and NaV1.6 in neocortical pyramidal cell excitability. eLife 14. Ed. by J. R. Huguenard. Publisher: eLife Sciences Publications, Ltd, RP105696. 10.7554/eLife.105696.

75. Radman, T., Ramos, R. L., Brumberg, J. C., and Bikson, M. (2009). Role of Cortical Cell Type and Morphology in Sub- and Suprathreshold Uniform Electric Field Stimulation. Brain stimulation 2, 215–228. 10.1016/j.brs.2009.03.007.

76. Kaczmarek, L. K. and Zhang, Y. (2017). Kv3 Channels: Enablers of Rapid Firing, Neurotransmitter Release, and Neuronal Endurance. Physiological Reviews 97. Publisher: American Physiological Society, 1431–1468. 10.1152/physrev.00002.2017.

77. Bikson, M., Inoue, M., Akiyama, H., Deans, J. K., Fox, J. E., Miyakawa, H., and Jefferys, J. G. R. (2004). Effects of uniform extracellular DC electric fields on excitability in rat hippocampal slices in vitro. eng. The Journal of Physiology 557, 175–190. 10.1113/jphysiol.2003.055772.

78. DeFelipe, J. and Fariñas, I. (1992). The pyramidal neuron of the cerebral cortex: morphological and chemical characteristics of the synaptic inputs. eng. Progress in Neurobiology 39, 563–607. 10.1016/0301-0082(92)90015-7.

79. Kawaguchi, Y. and Kubota, Y. (1997). GABAergic cell subtypes and their synaptic connections in rat frontal cortex. eng. Cerebral Cortex (New York, N.Y.: 1991) 7, 476–486. 10.1093/cercor/7.6.476.

80. Tremblay, R., Lee, S., and Rudy, B. (2016). GABAergic interneurons in the neocortex: From cellular properties to circuits. Neuron 91, 260–292. 10.1016/j.neuron.2016.06.033.

81. Martina, M., Schultz, J. H., Ehmke, H., Monyer, H., and Jonas, P. (1998). Functional and Molecular Differences between Voltage-Gated K+ Channels of Fast-Spiking Interneurons and Pyramidal Neurons of Rat Hippocampus. en. Journal of Neuroscience 18. Publisher: Society for Neuroscience Section: ARTICLE, 8111–8125. 10.1523/JNEUROSCI.18-20-08111.1998.

82. Gouwens, N. W., Sorensen, S. A., Baftizadeh, F., Budzillo, A., Lee, B. R., Jarsky, T., Alfiler, L., Baker, K., Barkan, E., Berry, K., et al. (2020). Integrated Morphoelectric and Transcriptomic Classification of Cortical GABAergic Cells. Cell 183, 935–953.e19. 10.1016/j.cell.2020.09.057.

83. Lee, S. Y., Kozalakis, K., Baftizadeh, F., Campagnola, L., Jarsky, T., Koch, C., and Anastassiou, C. A. (2024a). Cell-class-specific electric field entrainment of neural activity. eng. Neuron 112, 2614–2630.e5. 10.1016/j.neuron.2024.05.009.

84. Lee, J.-I., Werginz, P., Kameneva, T., Im, M., and Fried, S. I. (2024b). Membrane depolarization mediates both the inhibition of neural activity and cell-type-differences in response to high-frequency stimulation. en. Communications Biology 7. Publisher: Nature Publishing Group, 734. 10.1038/s42003-024-06359-3.

85. Penfield, W. and Boldrey, E. (1937). SOMATIC MOTOR AND SENSORY REPRESENTATION IN THE CEREBRAL CORTEX OF MAN AS STUDIED BY ELECTRICAL STIMULATION1. Brain 60, 389–443. 10.1093/brain/60.4.389.

86. Tehovnik, E. J. and Slocum, W. M. (2007). Phosphene Induction by Microstimulation of Macaque V1. Brain research reviews 53, 337–343. 10.1016/j.brainresrev.2006.11.001.

87. Hubel, D. H. and Wiesel, T. N. (1962). Receptive fields, binocular interaction and functional architecture in the cat’s visual cortex. The Journal of Physiology 160, 106–154.2. 10.1113/jphysiol.1962.sp006837.

88. Grinvald, A., Lieke, E., Frostig, R. D., Gilbert, C. D., and Wiesel, T. N. (1986). Functional architecture of cortex revealed by optical imaging of intrinsic signals. eng. Nature 324, 361–364. 10.1038/324361a0.

89. Livingstone, M. and Hubel, D. (1988). Segregation of form, color, movement, and depth: anatomy, physiology, and perception. eng. Science (New York, N.Y.) 240, 740–749. 10.1126/science.3283936.

90. Mountcastle, V. B. (1997). The columnar organization of the neocortex. Brain 120, 701– 722. 10.1093/brain/120.4.701.

91. Mazer, J. A., Vinje, W. E., McDermott, J., Schiller, P. H., and Gallant, J. L. (2002). Spatial frequency and orientation tuning dynamics in area V1. Proceedings of the National Academy of Sciences of the United States of America 99, 1645–1650. 10.1073/pnas.022638499.

92. Ringach, D. L., Shapley, R. M., and Hawken, M. J. (2002). Orientation selectivity in macaque V1: diversity and laminar dependence. eng. The Journal of Neuroscience: The Official Journal of the Society for Neuroscience 22, 5639–5651. 10.1523/JNEUROSCI.22-13-05639.2002.

93. Papale, P., Wang, F., Self, M. W., and Roelfsema, P. R. (2025). An extensive dataset of spiking activity to reveal the syntax of the ventral stream. english. Neuron 113. Publisher: Elsevier, 539–553.e5. 10.1016/j.neuron.2024.12.003.

94. Kay, K. N., Naselaris, T., Prenger, R. J., and Gallant, J. L. (2008). Identifying natural images from human brain activity. en. Nature 452. Publisher: Nature Publishing Group, 352–355. 10.1038/nature06713.

95. Mejias, J. F., Murray, J. D., Kennedy, H., and Wang, X.-J. (2016). Feedforward and feedback frequency-dependent interactions in a large-scale laminar network of the primate cortex. Science Advances 2. Publisher: American Association for the Advancement of Science, e1601335. 10.1126/sciadv.1601335.

96. Siegle, J. H., Jia, X., Durand, S., Gale, S., Bennett, C., Graddis, N., Heller, G., Ramirez, T. K., Choi, H., Luviano, J. A., et al. (2021). Survey of spiking in the mouse visual system reveals functional hierarchy. en. Nature 592. Publisher: Nature Publishing Group, 86–92. 10.1038/s41586-020-03171-x.

97. Doty, R. W. (1965). Conditioned reflexes elicited by electrical stimulation of the brain in macaques. Journal of Neurophysiology 28. Publisher: American Physiological Society, 623–640. 10.1152/jn.1965.28.4.623.

98. Doty, R. W. (1969). Electrical stimulation of the brain in behavioral context. eng. Annual Review of Psychology 20, 289–320. 10.1146/annurev.ps.20.020169.001445.

99. Rebesco, J. M., Stevenson, I. H., Körding, K. P., Solla, S. A., and Miller, L. E. (2010). Rewiring Neural Interactions by Micro-Stimulation. Frontiers in Systems Neuroscience 4, 39. 10.3389/fnsys.2010.00039.

100. Ni, A. M. and Maunsell, J. H. R. (2010). Microstimulation Reveals Limits in Detecting Different Signals from a Local Cortical Region. english. Current Biology 20. Publisher: Elsevier, 824–828. 10.1016/j.cub.2010.02.065.

101. Houweling, A. R. and Brecht, M. (2008). Behavioural report of single neuron stimulation in somatosensory cortex. en. Nature 451. Publisher: Nature Publishing Group, 65–68. 10.1038/nature06447.

102. Wolfe, J., Houweling, A. R., and Brecht, M. (2010). Sparse and powerful cortical spikes. eng. Current Opinion in Neurobiology 20, 306–312. 10.1016/j.conb.2010.03.006.

103. Histed, M. H., Ni, A. M., and Maunsell, J. H. R. (2013). Insights into cortical mechanisms of behavior from microstimulation experiments. Progress in Neurobiology. Conversion of Sensory Signals into Perceptions, Memories and Decisions 103, 115–130. 10.1016/j.pneurobio.2012.01.006.

104. Shah, N. P., Phillips, A., Madugula, S., Lotlikar, A., Gogliettino, A. R., Hays, M. R., Grosberg, L., Brown, J., Dusi, A., Tandon, P., et al. (2024). Precise control of neural activity using dynamically optimized electrical stimulation. eLife 13. Ed. by M. Beyeler, J. I. Gold, and M. Shivdasani. Publisher: eLife Sciences Publications, Ltd, e83424. 10.7554/eLife.83424.

105. Moreno, J. S., Garcia, N., and Denman, D. J. (2025). Separable neural population representations are constructed from mixed single neuron selectivity in the mouse early visual system. eng. bioRxiv: The Preprint Server for Biology, 2025.08.25.672226. 10.1101/2025.08.25.672226.

106. Siegle, J. H., López, A. C., Patel, Y. A., Abramov, K., Ohayon, S., and Voigts, J. (2017). Open Ephys: an open-source, plugin-based platform for multichannel electrophysiology. en. Journal of Neural Engineering 14. Publisher: IOP Publishing, 045003. 10.1088/1741-2552/aa5eea.

107. Pachitariu, M., Sridhar, S., Pennington, J., and Stringer, C. (2024). Spike sorting with Kilosort4. en. Nature Methods 21. Publisher: Nature Publishing Group, 914–921. 10.1038/s41592-024-02232-7.

108. Birman, D., Yang, K. J., West, S. J., Karsh, B., Browning, Y., Laboratory, t. I. B., Siegle, J. H., and Steinmetz, N. A. (2023). Pinpoint: trajectory planning for multi-probe electrophysiology and injections in an interactive web-based 3D environment. en. eLife 12. Publisher: eLife Sciences Publications Limited. 10.7554/eLife.91662.1.

109. Claudi, F., Tyson, A. L., Petrucco, L., Margrie, T. W., Portugues, R., and Branco, T. (2021). Visualizing anatomically registered data with brainrender. eLife 10. Ed. by M. W. Mathis, K. M. Wassum, and J. Nunez-Iglesias. Publisher: eLife Sciences Publications, Ltd, e65751. 10.7554/eLife.65751.

110. Montijn, J. S., Seignette, K., Howlett, M. H., Cazemier, J. L., Kamermans, M., Levelt, C. N., and Heimel, J. A. (2021). A parameter-free statistical test for neuronal responsiveness. eLife 10. Ed. by M. C. van Rossum, J. I. Gold, and J. D. Victor. Publisher: eLife Sciences Publications, Ltd, e71969. 10.7554/eLife.71969.

111. Rübel, O., Tritt, A., Ly, R., Dichter, B. K., Ghosh, S., Niu, L., Baker, P., Soltesz, I., Ng, L., Svoboda, K., et al. (2022). The Neurodata Without Borders ecosystem for neurophysiological data science. eLife 11. Ed. by L. L. Colgin and S. P. Jadhav. Publisher: eLife Sciences Publications, Ltd, e78362. 10.7554/eLife.78362.

